# Machine learning reveals limited contribution of trans-only encoded variants to the HLA-DQ immunopeptidome by accurate and comprehensive HLA-DQ antigen presentation prediction

**DOI:** 10.1101/2022.09.14.507934

**Authors:** Jonas Birkelund Nilsson, Saghar Kaabinejadian, Hooman Yari, Bjoern Peters, Carolina Barra, Loren Gragert, William Hildebrand, Morten Nielsen

**Affiliations:** Department of Health Technology, Technical University of Denmark, DK-2800 Lyngby, Denmark; Pure MHC, LLC., Oklahoma City, OK, United States; Department of Pathology and Laboratory Medicine, Tulane University School of Medicine, New Orleans, LA 70112, USA; Center for Infectious Disease and Vaccine Research, La Jolla Institute for Immunology, La Jolla, CA 92037, California, USA; Department of Microbiology and Immunology, University of Oklahoma Health Sciences Center, Oklahoma City, OK, United States

## Abstract

HLA class II antigen presentation is key for controlling and triggering T cell immune responses. HLA-DQ molecules, which are believed to play a major role in autoimmune diseases, are heterodimers that can be formed as both cis and trans variants depending on whether the α- and β-chains are encoded on the same (cis) or opposite (trans) chromosomes. So far, limited progress has been made for predicting HLA-DQ antigen presentation. In addition, the contribution of trans-only variants (i.e. variants not observed in the population as cis) in shaping the HLA-DQ immunopeptidome remains largely unresolved. Here, we seek to address these issues by integrating state-of-the-art immunoinformatics data mining models with large volumes of high-quality HLA-DQ specific MS-immunopeptidomics data. The analysis demonstrated a highly improved predictive power and molecular coverage for models trained including these novel HLA-DQ data. More importantly, investigating the role of trans-only HLA-DQ variants revealed a limited to no contribution to the overall HLA-DQ immunopeptidome. In conclusion, this study has furthered our understanding of HLA-DQ specificities and has for the first time cast light on the relative role of cis versus trans-only HLA-DQ variants in the HLA class II antigen presentation space. The developed method, NetMHCIIpan-4.2, is available at https://services.healthtech.dtu.dk/services/NetMHCIIpan-4.2.

## Introduction

Major histocompatibility complex class II molecules (MHC class II) are expressed on the surface of professional antigen presenting cells such as B cells, dendritic cells (DCs), and monocytes/macrophages^1^. These molecules, which are designed to bind and present fragments of the exogenous proteins to T-helper cells, are heterodimers consisting of α- and β-chains which together form the peptide-binding cleft.

In humans, HLA (human leukocyte antigen) class II is encoded by three different loci (HLA-DR, -DQ, and -DP). These HLA genes have numerous allelic variants with polymorphisms that are mainly clustered around the peptide-binding groove, resulting in a wide range of distinct peptide-binding specificities^2^. In many autoimmune diseases, HLA class II genes are major genetic susceptibility factors^1,3^ that play a central role in the pathogenesis of these conditions by presenting antigenic peptides to CD4+ T cells.

Several studies have explored the importance of HLA-DR and DQ at haplotype and genotype levels among type 1 diabetes (T1D) patients^3^. These genetic and functional studies have indicated that both HLA-DR and DQ alleles are associated with the risk of T1D^3,4^. In addition, the associated DR-DQ haplotypes demonstrate a risk hierarchy, ranging from highly predisposing to highly protective^4^. Interestingly, more recently it was demonstrated that HLA-DR, which generally plays the primary role in autoimmune diseases, has an important but secondary role to the HLA-DQ locus in T1D^5^.

Autoimmune disorders like T1D in addition to other conditions such as Celiac disease, where a direct and exceptionally strong association for HLA-DQ has been established^6^, thus necessitate a more thorough and systematic characterization of antigen presentation by HLA-DQ molecules to enable study of their function. Even though the field is moving forward rapidly^7^, so far peptide binding motifs of only a limited number of HLA-DQ molecules have been exhaustively studied^8–10^. One reason for this is that HLA-DQ molecules are more complex to study experimentally. For instance, because of the monomorphic nature of the α-chain in HLA-DR, the polymorphic variations are only provided by the β-chain^11^. In HLA-DQ, both α- and β-chains contribute to polymorphic variations. However, evidence suggests that not every α- and β-chain pairing will result in a stable heterodimer due to key structural requirements on the α and β dimerization interface^11,12^. For example, DQA1*01 has only been detected to form stable heterodimers with DQB1*05 and 06 alleles. Likewise, the DQA1*02, 03, 04, 05, and 06 alleles form stable heterodimers only with the DQB1*02, 03, and 04^12–14^.

In addition, studying the function of HLA-DQ alleles is challenging because of the extensive linkage disequilibrium between HLA-DR and HLA-DQ within the HLA class II region, making it difficult to differentiate the role of individual HLA-DQ alleles from the associated HLA-DR molecules^3,11^.

Finally, unique *cis* and *trans* encoded DQ molecules can occur where α- and β-chains that pair to form the heterodimer are encoded by the same (*cis*) or opposite (*trans*) chromosomes, making the study of these molecules even further complicated. While the majority of the current knowledge on HLA-DQ molecules comes from cis encoded variants, the surface expression and function of a small number of trans encoded DQ variants have been confirmed^11,15^. Here, it is important to emphasize that these functional trans molecules have also been observed to be functional as the corresponding cis-encoded variant. Therefore, it is generally believed that alleles of DQα- and DQβ-chains pair up primarily in cis rather than in trans variants^16,17^.

Hereafter, we refer to all stable DQα- and β-chain combinations mentioned above as cis, and the rest which includes any combination that has not been detected or reported as cis encoded will be referred to as “trans-only”.

In recent years, the information related to *cis*-encoded HLA-DQ variants has been greatly expanded due to large volumes of HLA sequence data becoming available^13^. Here, the assumption is that all observed DQ haplotypes, by natural selection, are able to form stable and functional cis and trans-encoded molecules. However, the role of trans-only encoded variants in antigen presentation and their contribution in shaping and complementing the HLA-DQ immunopeptidome has remained largely unresolved.

Given the critical role of HLA class II antigen presentation in the control and shaping of the adaptive immune response, great efforts have been dedicated to the development of prediction models capable of predicting this event (reviewed in Nielsen et al. 2020^18^). Current state-of-the-art prediction methods include NetMHCIIpan^19^, a pan-specific method allowing for prediction of antigen presentation for any HLA class II molecule with known protein sequence. For HLA-DQ and DP heterodimers, this means that sequence information about both the α- and β-chains is required in order to make predictions.

Originally, in vitro peptide-HLA binding affinity (BA) assays have been used to generate data to characterize the motifs of HLA class II molecules^2^, and development of different machine-learning prediction models to identify the rules of peptide–HLA binding^20,21^. However, experimental results indicate binding affinity (BA) to be a relatively weak correlate of antigen processing and presentation by HLA molecules^22^. In addition, multiple studies have demonstrated that the performance of the HLA-class II peptide-binding prediction models improve significantly when trained with immunopeptidome data acquired by liquid chromatography coupled with mass spectrometry (LC-MS/MS)^2,20,23,24^. Generally, in an HLA class II immunopeptidome eluted ligand (EL) assay, HLA molecules are affinity purified from lysed antigen presenting cells (APCs) using HLA specific monoclonal antibodies. The HLA molecules are next denatured and peptide ligands are isolated and sequenced via LC-MS/MS^25,26^. The result of such an assay is a list of peptide sequences restricted to at least one of the HLA class II molecules expressed by the interrogated cell line. EL data has a major advantage over BA data as they contain signals from different steps of HLA class II antigen presentation, such as antigen digestion, HLA loading of ligands, and transport to the cell surface^27–29^.

HLA class II binding predictions have been widely used to identify epitope candidates in infectious, cancer and autoimmune diseases^30^. The majority of prediction algorithms for HLA class II have so far been focused on HLA-DR molecules due to the large data availability for those. However, in the context of HLA-DQ, both pairing of synthetic α- and β-chains in order to perform binding affinity experiments, and generation of large EL datasets have proven to be challenging. The latter mostly due to lack of application of HLA-DQ specific antibodies in large scale MS-immunopeptidomics experiments resulting in limited yield in the HLA-DQ purification process.

In recent years, proteomics and peptide analysis by mass spectrometry (MS) has made huge progress, due to cutting edge technology and increased sensitivity of the instruments along with advanced software platforms and algorithms that support peptide identification and quantification. These advancements, along with the use of a highly specific HLA-DQ antibody, have enabled us to characterize, in a single assay, thousands of peptides which naturally bind the HLA-DQ molecules and generate stable peptide-HLA complexes that are transported to the cell surface to be presented to immune cells. Here, we have applied this setup to generate a large set of peptides presented by a group of HLA-DQ molecules frequent in the worldwide population from a panel of homozygous B lymphoblastoid cell lines. These large data sets were directly submitted to bioinformatic motif identification and machine learning pipelines to define the motifs and uncover the rules governing the processing and presentation of peptides in a biological context. Further, this study allowed us to move towards resolving the challenge of cis versus trans formation of functional HLA-DQ heterodimers and determine the role of trans-only variants in shaping the HLA-DQ immunopeptidome. The extensive insight into the peptide-binding characteristics of the investigated HLA-DQ molecules provided by this study will facilitate better understanding of HLA-DQ disease association and discovery of novel therapeutic targets.

## Materials and methods

### Cell lines and antibody

Homozygous B lymphoblastoid cell lines (BLCL) were obtained from the International Histocompatibility Working Group (IHWG) Cell and DNA bank housed at the Fred Hutchinson Cancer Research Center, Seattle, WA (http://www.ihwg.org). A group of 16 cell lines expressing the high frequency HLA-DQ alleles were selected for the study (supplementary table 1). To guarantee intact class II processing and presentation machinery and to ensure that the total HLA-DQ expression represents the physiological level, use of engineered cells was avoided.

The cells were grown in high density cultures in roller bottles in complete RPMI medium (Gibco) supplemented with 15% fetal bovine serum (FBS; Gibco/Invitrogen Corp) and 1% 100 mM sodium pyruvate (Gibco). Cells were harvested from the suspension, washed with PBS and spun down at 4C for 10 minutes. The cell pellets were immediately frozen in LN2 and stored at -80 until downstream processing^23^. All cell lines were subjected to high-resolution HLA typing (HLA-A, -B, -C, DRB1,3, 4, 5, DP and DQ) immediately upon receipt and growth in our laboratory, for authentication prior to large scale culture and data collection. The anti-human HLA-DQ specific monoclonal antibody was produced in house from a hybridoma cell line (clone SPVL3) and used for affinity purification of total HLA DQ from the BLCLs.

### Isolation and purification of HLA-DQ bound peptides

HLA-DQ molecules were purified from the cells by affinity chromatography using the anti-human HLA-DQ specific antibody (clone SPVL3) coupled to CNBr-activated Sepharose 4 Fast Flow (Amersham Pharmacia Biotech, Orsay, France) as described previously^23^. Briefly, frozen cell pellets were pulverized using Retsch Mixer Mill MM400, resuspended in lysis buffer comprised of Tris pH 8.0 (50 mM), Igepal, 0.5%, NaCl (150 mM) and complete protease inhibitor cocktail (Roche, Mannheim, Germany). Lysates were centrifuged in an Optima XPN-80 ultracentrifuge (Beckman Coulter, IN, USA) and filtered supernatants were loaded on immunoaffinity columns overnight at 4C. Columns were washed sequentially with a series of wash buffers^26^ and were eluted with 0.2 N acetic acid. The HLA was denatured, and the peptides were isolated by adding glacial acetic acid and heat. The mixture of peptides and HLA-DQ was subjected to reverse phase high performance liquid chromatography (RP-HPLC).

### MS

#### Fractionation of the HLA/Peptide Mixture by RP-HPLC

RP-HPLC was used to reduce the complexity of the peptide mixture eluted from the affinity column. First, the eluate was dried under vacuum using a CentriVap concentrator (Labconco, Kansas City, Missouri, USA). The solid residue was dissolved in 10% acetic acid and fractionated using a Paradigm MG4 instrument (Michrom BioResources, Auburn, California, USA). An acetonitrile (ACN) gradient was run at pH 2 using a two-solvent system. Solvent A contained 2% ACN in water, and solvent B contained 5% water in ACN. Both solvent A and Solvent B contained 0.1% trifluoroacetic acid (TFA). The column was pre-equilibrated at 2% solvent B. Then the sample was loaded at a flow rate of 120 µl/min and a two-segment gradient was run at 160 µl/min flow rate as described in detail in Kaabinejadian et al. 2022^23^. Fractions were collected in 2 min intervals using a Gilson FC 203B fraction collector (Gilson, Middleton, Wisconsin, USA), and the ultra-violet (UV) absorption profile of the eluate was recorded at 215 nm wavelength.

#### Nano LC-MS/MS Analysis

Peptide-containing HPLC fractions were dried and resuspended in a solvent composed of 10% acetic acid, 2% ACN and iRT peptides (Biognosys, Schlieren, Switzerland) as internal standards. Fractions were applied individually to an Eksigent nanoLC 415 nanoscale RP-HPLC (AB Sciex, Framingham, Massachusetts, USA), including a 5-mm long, 350 µm internal diameter Chrom XP C18 trap column with 3 µm particles and 120 Å pores, and a 15-cm-long ChromXP C18 separation column (75 µm internal diameter) packed with the same medium (AB Sciex, Framingham, Massachusetts, USA). An ACN gradient was run at pH 2.5 using a two-solvent system. Solvent A was 0.1% formic acid in water, and solvent B was 0.1% formic acid in 95% ACN in water. The column was pre-equilibrated at 2% solvent B. Samples were loaded at 5 μL/min flow rate onto the trap column and run through the separation column at 300 nL/min with two linear gradients: 10% to 40% B for 70 minutes, followed by 40% to 80% B for 7 minutes.

The column effluent was ionized using the nanospray III ion source of an AB Sciex TripleTOF 5600 quadruple time-of-flight mass spectrometer (AB Sciex, Framingham, MA, USA) with the source voltage set to 2,400 V. Information-dependent analysis (IDA) method was used for data acquisition^23^. PeakView Software version 1.2.0.3 (AB Sciex, Framingham, MA, USA) was used for data visualization.

### Peptide Data Analysis

Peptide sequences were identified using PEAKS Studio 10.5 software (Bioinformatics Solutions, Waterloo, Canada). A database composed of SwissProt Homo sapiens (taxon identifier 9606) and iRT peptide sequences was used as the reference for database search. Variable post-translational modifications (PTM) including acetylation, deamination, pyroglutamate formation, oxidation, sodium adducts, phosphorylation, and cysteinylation were included in database search. Identified peptides were further filtered at a false discovery rate (FDR) of 1% using PEAKS decoy-fusion algorithm.

#### Immunopeptidome Data

The immunopeptidome data consist of MS-eluted ligand (EL) and binding affinity (BA) data from the earlier NetMHCIIpan-4.1 combined with the EL data generated specifically for this study (see above). The novel MS-immunopeptidome data set covers 14 different HLA-DQ molecules obtained from 16 homozygous BLCLs. This data was filtered as described earlier^23^ to exclude potential HLA class I and other co-immunoprecipitated contaminants, resulting in a list of peptides of length 12-21.

The EL data were mapped to the human reference source proteome to define source protein context. Peptides with no identical reference match were excluded, resulting in ∼4% of peptides being discarded. Finally, the EL data were enriched in a per sample-id manner with random natural peptides assigned as negatives. This enrichment was done by sampling peptides of 12-21 amino acids in length in a uniform manner in an amount equal to five times the number of peptides for the most prevalent length in the positive data for the given sample.

Our final novel data set consists of 39,334 positive and 369,313 negative peptides covering 14 unique HLA-DQ molecules. This data set is available in supplementary table 2. Merging the novel EL data with the earlier NetMHCIIpan-4.1 data (expanded to include peptides 12 amino acids in length), the complete EL data consists of 480,845 positive and 4,910,165 negative data points from 177 samples/cell lines, and the BA data consist of 129,110 data points.

The data was partitioned into five subsets for cross-validated method training and evaluation using the common-motif approach described earlier^31^ merging EL and BA data ensuring that peptides sharing an identical overlap of 9 or more consecutive amino acids were placed in the same subset.

## Methods

### Model training

Models were trained using the NNAlign_MA machine learning framework^32^ in a manner similar to that described earlier for NetMHCIIpan-4.0^2^. That is, the complete model consists of an ensemble of 100 neural networks of two different architectures both with one hidden layer and either 40 or 60 hidden neurons, with 10 random weight initializations for each of the 5 cross-validation folds (2 architectures, 10 seeds, and 5 folds). All models were trained using backpropagation with stochastic gradient descent, for 300 epochs, without early stopping, and a constant learning rate of 0.05. Only single allele (SA) data were included in the training for a burn-in period of 20 epochs.

Subsequent training cycles included multi-allele (MA) data. Two main models were trained, one including the original NetMHCIIpan-4.1 data and one including the novel HLA-DQ data. Furthermore, an additional model was trained with the novel data using peptide context encoding. Here, context was defined in both the peptide’s N- and C-terminal as three residues from the source protein flanking the peptide, along with three starting residues from the peptide, all concatenated into a 12-mer amino acid sequence. For further details refer to Barra et al. 2018^27^.

### Performance evaluation and MHC restriction deconvolution

HLA annotation for MA datasets is based on which HLA molecule expressed in a given cell line has the highest prediction score for a given ligand. To balance the differences in the prediction score distributions between HLAs, percentile normalized prediction scores for each molecule were generated by ranking against a distribution of prediction scores of random natural peptides as described earlier^19,33^.

Performance was evaluated on the concatenated cross-validation test set predictions using three separate metrics, namely AUC (Area Under the ROC Curve), AUC 0.1 (Area Under the ROC Curve integrated up to a False Positive Rate of 10%) and Positive Predictive Value (PPV). Each metric was calculated in a per-HLA manner from the “raw” prediction scores after HLA annotation. Further, the PPV was calculated as the fraction of true positives in the top N predictions, where N is the number of ligands assigned to a given HLA molecule. For the per-HLA performance evaluation, only HLA molecules with at least 10 positive peptides in both models were included in the performance evaluation, to ensure a level of certainty in the calculated performance metrics.

### Statistical tests

Statistical tests in the performance evaluation were performed using one-tailed binomial tests excluding ties, with a significance level of 0.05. The alternative hypothesis in these tests is thus that the model trained with the novel data is more likely to perform better on a given HLA molecule than the model trained without.

### Consistency correlation matrix analysis

In order to assess the novel DQ data’s impact on NNAlign_MA’s motif deconvolution, a consistency correlation matrix analysis was performed as described earlier^2^. To avoid potential MS co-immunoprecipitated contaminant peptides biasing this analysis, the union of identified trash peptides (i.e. positive peptides given a percentile rank greater than 20 in either of the two models) was removed. A position-specific scoring matrix (PSSM) was next generated for each molecule in each cell line based on the predicted peptide binding cores. Here, a minimum of 20 positive peptides was required in order for a PSSM to be generated. Then, for each pair of cell lines sharing a given molecule, the Pearson Correlation Coefficient (PCC) between the molecule’s PSSMs was calculated. The mean consistency value for a given molecule was then given as the average PCC over each unique cell line pair (excluding self-correlations). This metric thus indicates how consistent the identified binding motifs are across different datasets for each HLA class II molecule.

### Similarity distance measure

Distance between two HLA class II molecules was estimated as described earlier^34^ from the pseudo-distance of the two molecules, i.e.

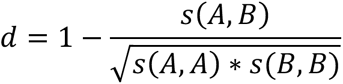

where s(X,Y) is the summed BLOSUM 50 similarity between the pseudo-sequences of molecule X and Y. Here, the pseudo-sequences are defined as described earlier from the set of 34 polymorphic residues within the HLA sequence concatenated into a continuous sequence, of which 15 and 19 residues derive from the α- and β-chain, respectively^35^.

### Estimation of prevalent stable HLA-DQ molecules

A list of HLA-DQ α- and β-chains forming prevalent stable HLA-DQ heterodimers was constructed by first obtaining lists of DQA1 and DQB1 alleles with annotated worldwide allele frequencies. This was done by querying the allelefrequencies.net database^36^ for high resolution alleles in populations of size 100 and above. Next, worldwide allele frequencies were obtained as population size weighted averages capping the maximum population size to 1000. Finally, a list of prevalent HLA-DQ molecules was constructed by pairing all α and β combinations following the restrictions outlined in table 1, only including molecules with a combined allele frequency greater than 0.00005. This resulted in a list of 154 HLA-DQ molecules.

**Table 1:** List of DQA1-DQB1 haplotypes extracted from Creary et al. and Petersdorf et al.^13,14^. Due to the relatively small sample size for some populations included in the study by Creary et al., the reported alleles and haplotypes may not reflect all haplotypes observed in the entire populations. Therefore, in this table only low-resolution (2 digit) DQ haplotypes were included to define the observed DQA1-DQB1 haplotypes.

| DQA1* - DQB1* | DQA1* - DQB1* |
| --- | --- |
| DQA1*01-DQB1*05 | DQA1*04-DQB1*02 |
| DQA1*01-DQB1*06 | DQA1*04-DQB1*03 |
| DQA1*02-DQB1*02 | DQA1*04-DQB1*04 |
| DQA1*02-DQB1*03 | DQA1*05-DQB1*02 |
| DQA1*02-DQB1*04 | DQA1*05-DQB1*03 |
| DQA1*03-DQB1*02 | DQA1*05-DQB1*04 |
| DQA1*03-DQB1*03 | DQA1*06-DQB1*02 |
| DQA1*03-DQB1*04 | DQA1*06-DQB1*03 |
|  | DQA1*06-DQB1*04 |

### Estimation of worldwide haplotype frequencies

Worldwide HLA-DQ haplotype frequencies were estimated by querying the allelefrequencies.net database^36^ for high resolution DQ haplotypes in populations of size 100 and above, average across population as described above for HLA-DQ frequencies.

### HLA-DQ specificity trees

An HLA-DQ specificity tree was constructed by first reducing the list of 154 prevalent HLA-DQ molecules to the set of unique pseudo-sequences among the molecules. Then, each unique pseudo-sequence was mapped to a representative HLA-DQ molecule name. By default, a DQ molecule in the list of molecules covered by the training data was used to represent a pseudo-sequence when possible. Furthermore, all 14 DQ molecules in the novel data were used to represent their given pseudo-sequences. In other cases of multiple options for a given pseudo-sequence, the most prevalent DQ molecule in terms of global allelic frequency was chosen. The specificity tree was then calculated using the MHCCluster method^37^ and visualized using the Iroki phylogenetic tree viewer^38^.

A similar tree was constructed based on clustering of the DQ pseudo-sequences. This tree was calculated with ClustalW-2.1^39^ using its phylogenetic tree function, and again visualized using the Iroki tree viewer^38^.

### Independent benchmark

For our benchmark against MixMHC2pred-2.0^7^, an independent dataset was taken from Marcu et al.^40^, which consists of eluted ligand data from 17 donor samples. This data was enriched with random negative peptides similarly to the training data. To reduce bias, peptides which were present in the EL training data of our method were not included in the benchmark. This yielded a total of 165,650 positive and 2,952,352 negative peptides covering 71 HLA class II molecules.

Performance was evaluated on a per-sample basis in terms of AUC, AUC 0.1, and PPV. For both our method and MixMHC2pred, the HLA annotation for each peptide was based on the lowest reported percentile rank scores for the HLA molecules in the given sample. Furthermore, as MixMHC2pred recommends using its reported percentile rank scores for all analyses, its predictive performance was calculated using these rank scores.

## Results

For the study, immunopeptidome data for 14 different HLA-DQ molecules was obtained from 16 homozygous B Lymphoblastoid Cell Lines (BLCLs) using LC-MS/MS. By using a DQ-specific antibody during the affinity purification, we were able to obtain a large dataset highly enriched in DQ peptide ligands. An overview of the cell lines’ peptide counts, DQ HLA types and peptide length distributions is shown in figure 1. Overall, the data contains a total of 39,334 peptide ligands, with 14- and 15-mers being most prevalent. After enriching the novel data with random natural peptides assigned as negatives (see materials and methods), we combined it with the data used to train the NetMHCIIpan-4.1 prediction method, yielding a large dataset of eluted HLA class II ligands. From this, we set out to address three essential issues related to HLA-DQ, namely i) the relatively low predictive power of current prediction models for DQ molecules, ii) the contribution of trans-only encoded DQ variants to the DQ immunopeptidome, and iii) the overall coverage of the DQ specificity space of the current experimental data and developed in-silico prediction models.

**Figure 1:**
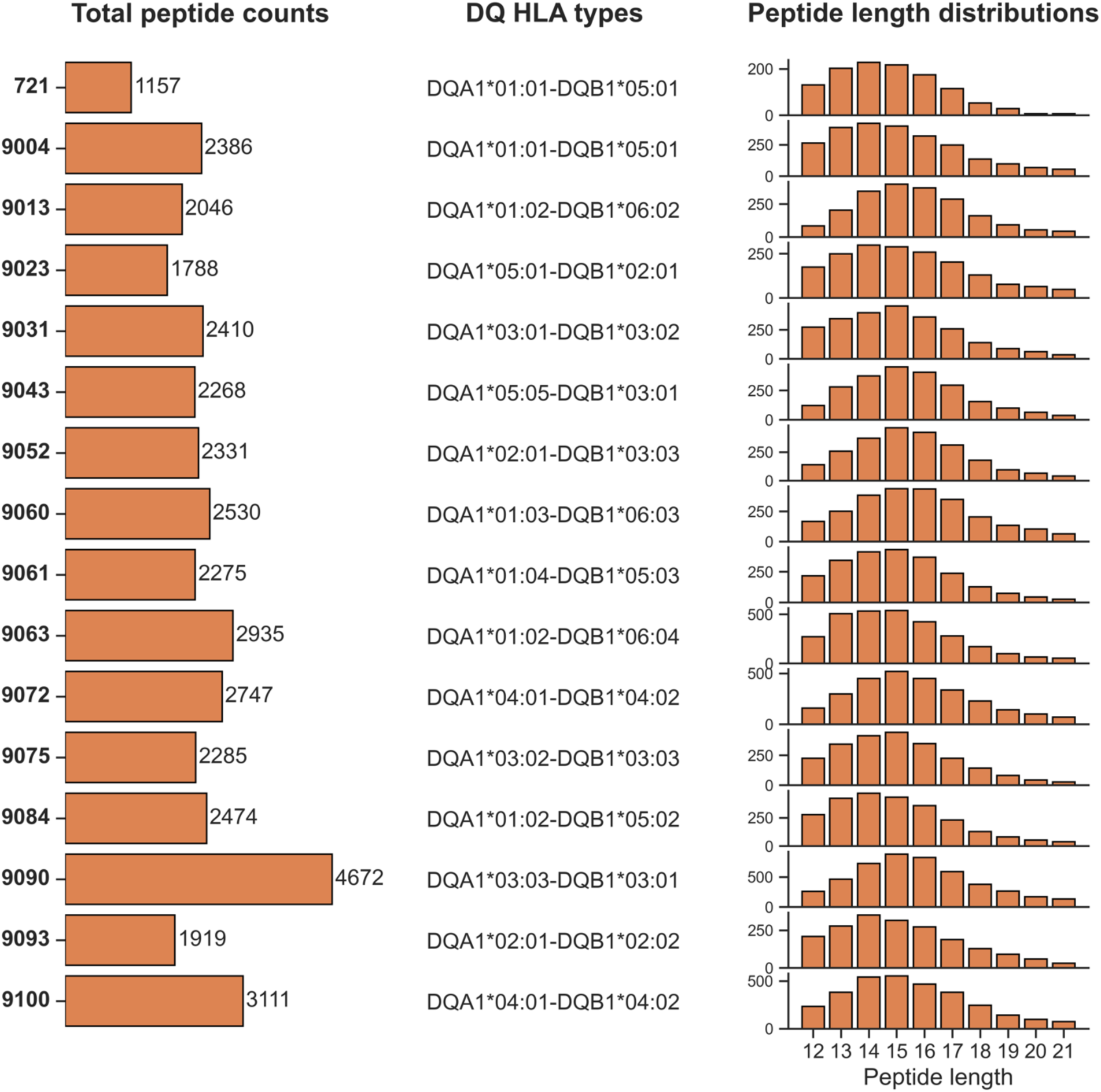
Overview of the novel immunopeptidomics data. Each row corresponds to a dataset from a given DQ-homozygous cell line. **Left panel**: Bar plot of overall peptide counts. The numbers on the left correspond to the cell line IDs. **Middle panel:** DQ HLA types of the cell lines. **Right panel**: Peptide length distributions.

### Impact of novel DQ data on predictive performance

To investigate the impact on the predictive power by integration of the novel DQ data, we employed the NNAlign_MA algorithm^32^ which is a highly powerful machine learning method for deconvoluting MS immunopeptidomics data. Two peptide antigen presentation prediction models were trained: one including the novel DQ affinity purified data (termed w_Saghar_DQ), and for direct comparison of the impact of the novel data one without (termed wo_Saghar_DQ). The models were then evaluated using cross-validation on a per-molecule basis within four different subsets of all the HLA class II molecules in the training data. These subsets are non-DQ molecules (NotDQ), all DQ molecules (DQ), DQ molecules present in the novel data (DQ_Saghar) and DQ molecules not present in the novel data (DQ_NotSaghar).

Figure 2 displays the result of this experiment and demonstrates that incorporation of the novel DQ data resulted in a significant performance gain for DQ as expected (p=0.011 for all metrics, one-tailed binomial test without ties). However, from these results it is apparent that the performance for DQ remains lower compared to that of non-DQ molecules. We assumed this to be a result of the DQ performance being calculated from a mix of both the novel data and the older NetMHCIIpan-4.1 training data. To demonstrate this, we evaluated the performance of the DQ_Saghar molecules limited to the novel data only. The result of this is shown in figure 3 and demonstrates that when focusing only on the novel data, the performance of DQ reaches a level comparable to that of non-DQ (median AUC values for DQ_Saghar and NotDQ are 0.9303 and 0.9312, respectively). This result is important as it suggests that the low performance earlier reported for DQ is at least in part imposed by a low quality and quantity of the earlier DQ data.

**Figure 2:**
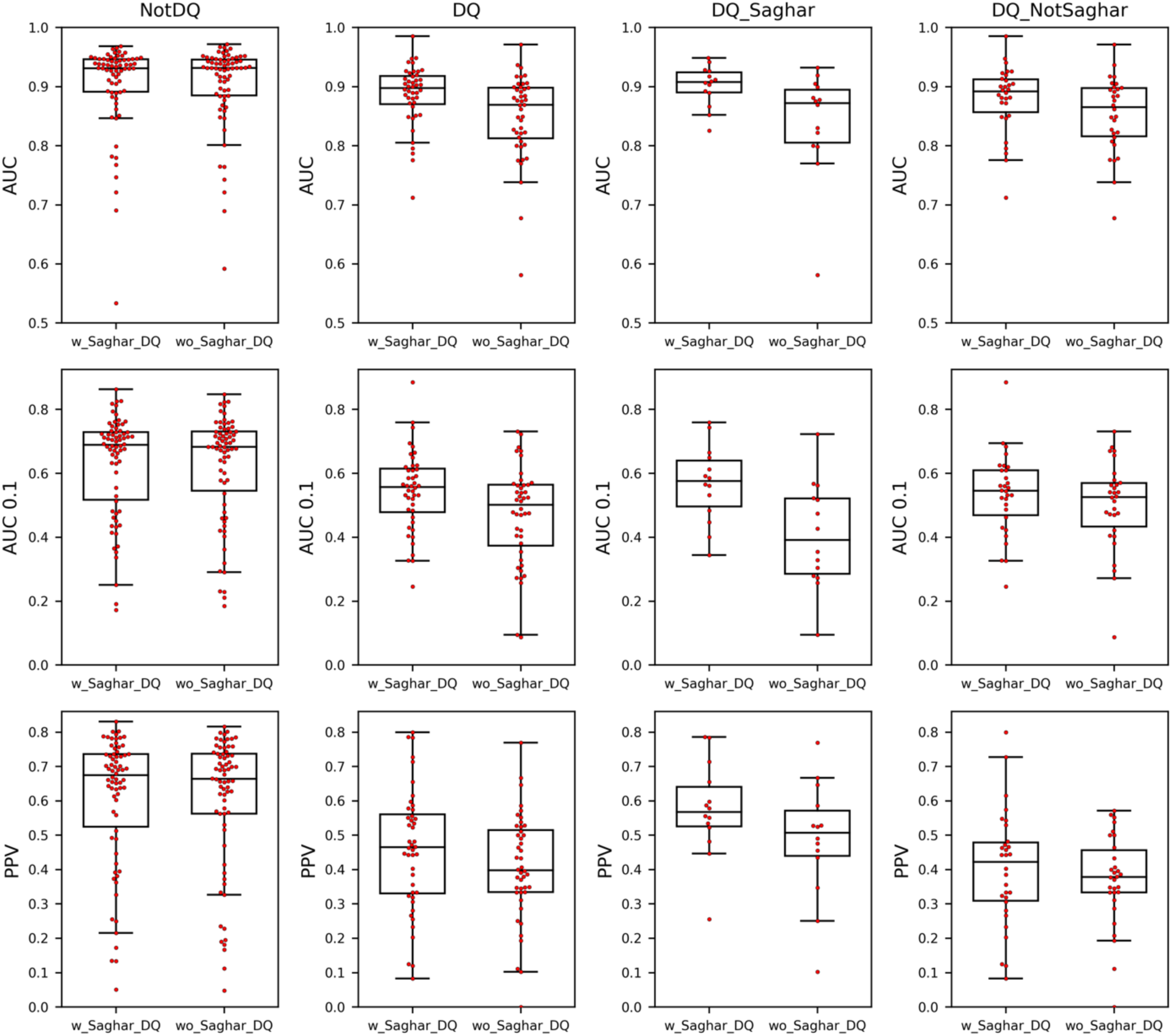
AUC, AUC 0.1 and PPV predictive performance for the models trained with (w_Saghar_DQ) and without (wo_Saghar_DQ) the novel data. Each point is the performance metric for a unique HLA class II molecule. For details on the performance metrics refer to materials and methods. The columns correspond to four different subsets of HLA molecules, namely all non-HLA-DQ molecules (NotDQ), all DQ molecules (DQ), DQ molecules in the novel data set (DQ_Saghar), and DQ molecules not present in the novel data (DQ_NotSaghar).

**Figure 3:**
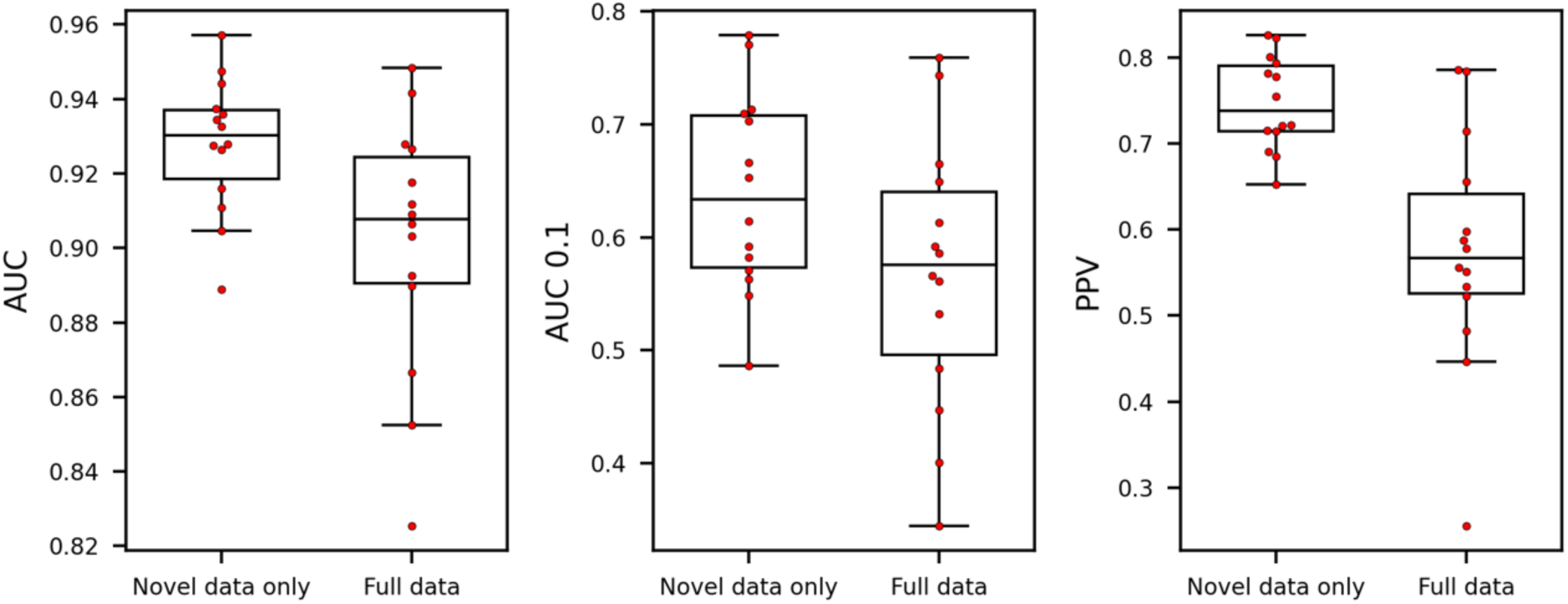
Performance of model trained including the novel data, evaluated on both the novel data alone, as well as the full dataset. Each point is the performance metric for an HLA-DQ molecule present in the novel data.

We next looked at the differences in peptides assigned to HLA-DQ molecules between the two methods across all samples. Here, we considered all peptides which were assigned to DQ with percentile rank less than 20 (i.e. as non-trash) in at least one of the methods^23^. Overall, the two methods share a high degree of overlap in the peptides assigned to DQ (60,959 annotations were shared by both models, 9309 annotations were unique for the method trained including the novel data and 4316 unique for the method trained without). This increased DQ coverage for the model trained including the novel data predominantly comes from peptides assigned to DR (and to some degree trash and DP) by the model trained without the novel data (see supplementary table 3 for an overview of the peptide migrations). This suggests that at least part of the improved predictive performance of the novel model originates from an improved motif deconvolution.

To further quantify this, we show the mean consistency value per HLA molecule in the four molecule subsets in supplementary figure 1. In short, position-specific scoring matrices were constructed for each molecule in a given cell line from the predicted binding cores in the individual positive peptides, and the consistency was quantified by the correlation of such matrices for the same molecule between different cell line data sets (for details refer to materials and methods). Based on this analysis, an overall improved consistency is observed for the model trained with the novel DQ data (p<0.02 in all cases except for the DQ_NotSaghar subset, one-tailed binomial test without ties). The consistency analysis for an example molecule contained in the novel data (DQA1*03:01-DQB1*03:02) is shown in supplementary figure 2, illustrating that in most cases the improved motif consistency is caused by an increased peptide count across samples (see supplementary tables 4 and 5).

Furthermore, HLA-DQ binding motifs obtained by motif deconvolution of the novel MS data were visualized, along with sequence motifs based on predicted binders, in supplementary figure 3. Here, the logos obtained by motif deconvolution are in most cases very similar when comparing the models trained with and without the novel data. However, the predicted sequence logos based on top scoring random natural peptides indicate that the model trained without the novel DQ data has failed to fully learn the correct binding motifs of all the novel DQ molecules, especially with respect to the P1 amino acid preferences. To quantify these results, correlations between the deconvoluted and predicted logos for each method were calculated (supplementary figure 4). This analysis showed significantly higher correlation for the method including the novel data (p=0.011, one-tailed binomial test without ties), indicating a highly consistent correspondence between the identified and predicted binding motifs.

Together, these observations demonstrate that incorporating the novel HLA-DQ data has allowed for an enriched identification of HLA-DQ peptide ligands, rescuing peptides otherwise assigned to alternative DR/DP molecules, resulting in improved motif deconvolution consistency and improved predictive power.

The above results were complemented by a comparison to a model trained including the novel data using peptide context encoding. In short, context encoding refers to a scenario where information from the regions flanking the peptide is extracted from the source protein sequence and included as additional input to the machine learning model. In line with what has been demonstrated earlier^2,27,32^, the results of this comparison (supplementary figure 5) demonstrated that the model trained including context significantly outperformed the model trained without context in all performance metrics and data subsets (the only exception being the DQ_NotSaghar subset). However, given that the main focus of the remaining part of the manuscript is to investigate motif deconvolution and the role of cis versus trans-only DQ α- and β-chain pairing in this context, we focus on the simpler model trained without context information from here on.

### Distribution of annotations to cis vs trans-only DQ molecules

In DQ-heterozygous cell lines, four possible α-β chain pairings can in principle be observed. For so-called cis-heterodimers, the α- and β-chain are expressed on the same chromosome and can thus be observed in haplotype sequencing. DQ molecules formed by pairing α- and β-chains between chromosomes are called trans-heterodimers. Some α-β pairings have not been observed as cis encoded (based on large HLA-haplotype sequencing population studies) and are thus here referred to as “trans-only” combinations. To assess the relative contribution of cis and trans-only DQ heterodimers in shaping the immunopeptidome, we investigated the distribution of peptides assigned to cis versus trans-only encoded DQ molecules across DQ-heterozygous datasets for the two models. Here, only datasets with at least 100 DQ-annotated peptides excluding trash in both methods were considered (for an overview of the datasets used in this analysis, refer to supplementary table 6). The proportion of DQ-annotated peptides assigned to each molecule was then calculated for each dataset containing that molecule. Finally, the mean per-dataset peptide fraction was reported for each DQ molecule, and the distribution of these means for molecules across four categories were then investigated. These categories are all cis variants, cis-SA (cis variants part of the DQ-SA training data), cis-MA (cis variants part of the DQ-MA training data), and trans-only variants.

The result of this analysis is shown in figure 4A for the two models and indicates that for the method including the novel data, trans-only molecules consistently cover a small proportion of the DQ annotations in each cell line. On the other hand, the cis molecules have generally high contribution, with the cis-SA molecules having the largest contribution as expected. However, the cis-MA molecules were also found to have significantly larger contribution than the trans-only molecules in the model including the novel data (p=0.005, two-sample t-test). Similar results were found when extending the cis-SA category to include cis-MA molecules with the same pseudo-sequence as a cis-SA molecule (supplementary figure 6). Further, an overall higher contribution of trans-only molecules to the DQ peptide annotations was observed for the model trained without the novel data (p=0.03, paired one-sided t-test). These results are striking, as they indicate that the motif deconvolution in the model including the novel data is not solely driven by the cis-SA molecules, but rather by an overall preference for cis-encoded variants compared to trans-only variants (see supplementary figures 7 and 8).

**Figure 4:**
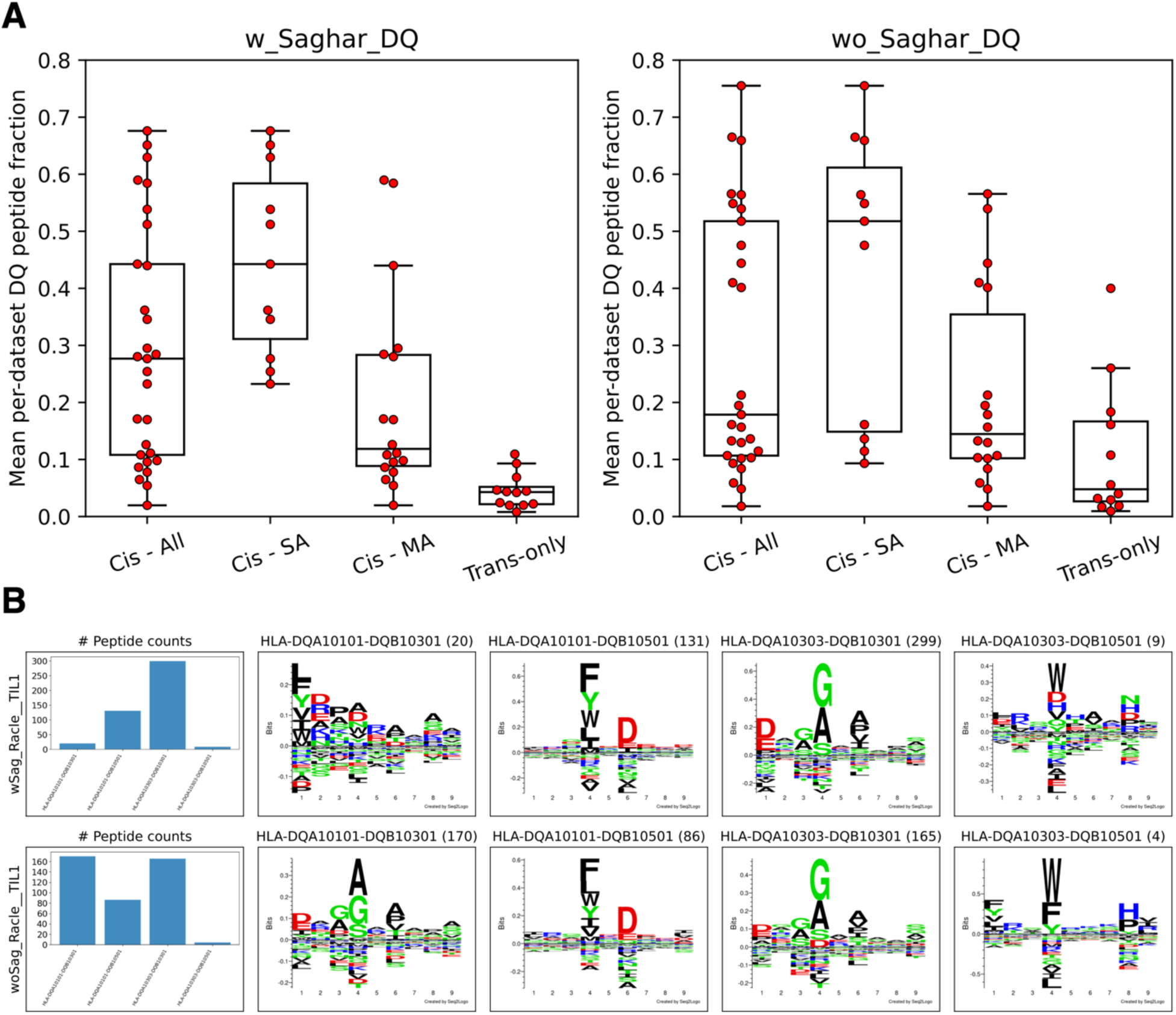
Contribution of cis and trans-only DQ variants in DQ-heterozygous datasets. **A:** Peptide-count contribution of cis and trans-only molecules in the methods with (w_Saghar_DQ) and without (wo_Saghar_DQ) the novel data. Each point shows the mean per-dataset peptide fraction for a given DQ molecule. The cis molecules are shown in three categories, namely all cis molecules (Cis - All), cis molecules found in the DQ-SA training data (Cis - SA), and cis molecules only found in the DQ-MA training data (Cis - MA). **B:** DQ motif deconvolution for the Racle TIL1 dataset. The rows correspond to the methods trained with (wSag) and without (woSag) the novel data, respectively. Peptide counts (excluding trash peptides) are displayed in parenthesis in the logo plot titles.

To further investigate this, the DQ motif deconvolution of the two models for the Racle TIL1 dataset is shown in figure 4B. Here, the model trained without the novel data assigns a large proportion of peptides (170 out of 425) to HLA-DQA1*01:01-DQB1*03:01, which is a trans-only molecule known to not form a stable heterodimer^12,13^. On the other hand, in the model trained with the novel data, almost no peptides are assigned to this molecule (20 out of 459). Instead, the peptides are assigned to the cis molecule HLA-DQA1*03:03-DQB1*03:01. Note, also, that for both models a very minor proportion of peptides are assigned to HLA-DQA1*03:03-DQB1*05:01, another trans-only heterodimer known to be unstable^12,13^. Overall, these results demonstrate that the model including the novel DQ data allows for proper motif deconvolution with limited assignment of peptides to trans-only HLA-DQ molecules. Further, the very low proportion of peptides assigned to trans-only molecules, combined with the overall increased HLA-DQ peptide volume and motif consistency of the model trained including the novel data, strongly suggests that trans-only HLA-DQ molecules are unstable, non-functional and have limited to no contribution to the total HLA-DQ immunopeptidome.

### Difference in peptide length distributions of DR and DQ

When we compared the length distribution of DQ peptide ligands in the novel data with HLA-DR restricted peptides that were purified from the same set of BLCLs^23^, it was revealed that the DQ ligands were in general shorter than the DR ligands (see supplementary figure 9). By comparing the per-molecule median peptide lengths for the two loci, a significant difference was found (p<0.03, two-sample t-test), with DR and DQ having average peptide length medians of 15.41 and 14.93, respectively. This analysis indicates that HLA-DQ molecules generally bind shorter peptides compared to HLA-DR. Moreover, in contrast to HLA-DQ alleles that are more consistent in their peptide length preferences, various HLA-DR molecules show subtle differences in their length preferences^23^. For example, HLA-DR*07:01, 09:01 and 14:01 show a preference for shorter peptides (14 mers) while the majority of DR alleles follow the common class II length preference (15 mer).

### Coverage of DQ

Next, we wanted to assess the number of DQ molecules present in the cross-validation predictions by each model which were properly covered (i.e. had a large number of peptides assigned during training), and hence where the models are expected to achieve accurate predictive power. The peptide count for a given DQ molecule was estimated as the accumulated sum of peptides from each cell line containing that molecule (excluding trash peptides). Here, only peptides annotated to DQ molecules in a given cell line corresponding to at least 5% of the total number of DQ peptides were included in its count (this was done to avoid including accumulation of low count noise). A given DQ molecule was then said to be covered if the summed peptide count over all cell lines was at least 100. This analysis resulted in 24 DQ molecules being covered by the model trained including the novel data, and 23 being covered when excluding these data. None of the 24 DQ molecules covered by the model including the novel data were found to be trans-only, whereas the model without the novel data covered two trans-only DQ molecules, namely HLA-DQA1*01:01-DQB1*03:01 (as described earlier) and HLA-DQA1*01:03-DQB1*03:02. Of the remaining 21 molecules, 20 were included in the molecules covered by the model trained with the novel data.

Given the different sets of molecules covered by the two methods, we wanted to estimate each method’s coverage when considering the entire DQ specificity space. As such, for each of the two methods, we investigated the proportion of 154 prevalent DQ molecules that had a distance of at most 0.025 to a molecule covered by the model (this set of molecules is here referred to as ‘extended coverage’). For details on how this distance was determined and how the list of prevalent DQ molecules was defined refer to materials and methods. The threshold of 0.025 was chosen based on the distance at which the model trained without the novel data could reach optimal performance on molecules not part of the method’s DQ-SA training data (see supplementary figure 10). Note, also, that 0.025 is a conservative distance threshold, and that we expect the model to maintain accuracy also for molecules falling beyond this value^35^.

From this analysis, a significant gain in extended coverage was found (p=0.027, chi-squared test), with the model including the novel data covering 94 out of 154 molecules, while the model without the novel data only covered 75 out of 154 molecules (see supplementary tables 7 and 8 for a list of covered and non-covered DQ molecules for the model trained including the novel data). When comparing the covered and non-covered molecules for the method including the novel data, the non-covered group had significantly lower worldwide haplotype frequency data as obtained from Allelefrequencies.net (for detail on how these frequencies were obtained refer to material and methods) compared to the covered group (average frequencies for the two groups were 0.0134 and 0.0025, p<0.03, student t-test). These results suggest that the non-covered DQ molecules are of limited importance seen from a population coverage perspective.

For visualizing the coverage of the DQ space, a specificity tree was constructed. Here, we used the list of 154 prevalent HLA-DQ molecules as the starting point. This list was first reduced to a set of 61 molecules with unique specificities (for details see methods) which were included in the subsequent analysis. Next, a specificity tree was constructed covering the 61 DQ molecules applying the MHCCluster method^37^. In short, the MHCCluster method estimates the similarity between two MHC molecules using the correlation between predicted binding values for a large set of random natural peptides. Figure 5 shows the resulting specificity tree along with predicted binding motifs for the 14 novel DQ molecules. The tree displays wide coverage of the DQ space, as all the novel molecules are spread more or less uniformly across the different branches of the tree, and all branches are covered by one or more DQ molecules in close distance to the DQ molecules covered by the training data. Moreover, a few subclusters of non-covered molecules were observed (highlighted by motifs in red frames), which were found to correspond almost one-to-one with the non-covered clusters in a phylogenetic tree of the DQ pseudo-sequences (see supplementary figure 11).

**Figure 5:**
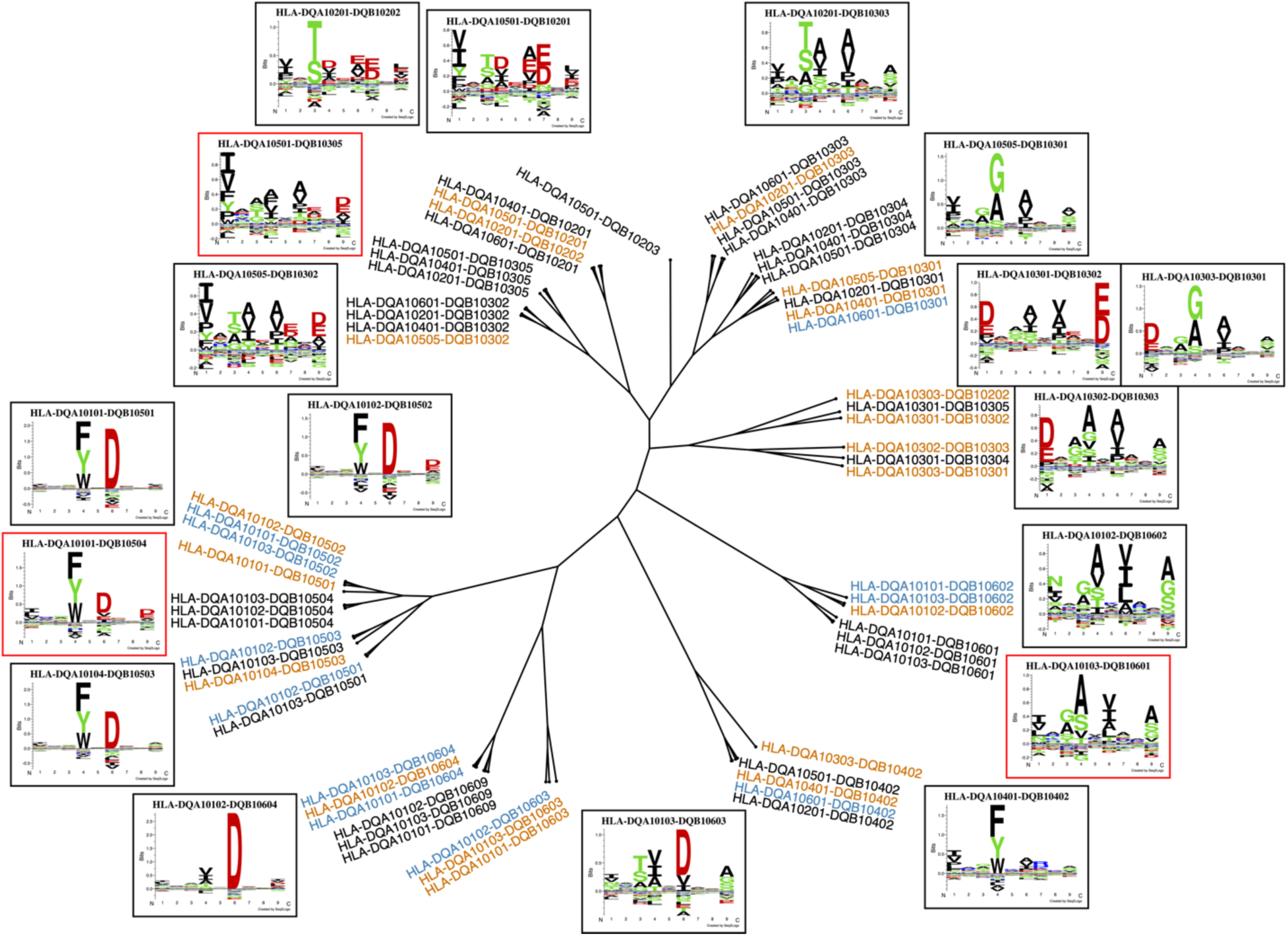
HLA-DQ specificity tree. The tree is based on 61 DQ molecules including the 14 molecules described by the novel data. Orange molecules are covered by the method including the novel data with at least 100 peptides, and blue molecules are within a distance 0.025 of an orange molecule. Black molecules are non-covered (i.e. have peptide count less than 100 and have distance greater than 0.025 to an orange molecule). Logos in black frames correspond to orange molecules. Logos in red frames correspond to molecules from branches with clusters of non-covered (black) molecules. The specificity tree was calculated from the pairwise similarities between the predictions scores for the DQ molecules for a set of 100,000 random natural 13-17mer peptides. Logos were constructed for the top 1% highest scoring binding cores for these 100,000 peptides.

### NetMHCIIpan-4.2

The model developed here including the novel DQ immunopeptidome data is made publicly available at https://services.healthtech.dtu.dk/service.php?NetMHCIIpan-4.2. The method allows for prediction of HLA antigen presentation to all HLA-DQ molecules, and prediction can be made with or without context encoding.

### Benchmark on independent DQ data

As a final showcase of our method’s motif deconvolution power for DQ, we benchmarked our method against MixMHC2pred-2.0, another HLA class II predictor which was recently published^7^. The benchmark data was taken from Marcu et al.^40^ and consists of eluted ligand data from 17 donor samples, which was enriched with random negative peptides.

Figure 6A shows the performance of the two methods per sample on the entire data, indicating that our method significantly outperforms MixMHC2pred on the independent dataset in all three metrics (p<0.05 in all metrics, one-tailed binomial test without ties). Furthermore, figure 6B shows the performance per sample restricted to the union of peptides annotated towards DQ by either method, once again showing significant performance gain in favor of NetMHCIIpan-4.2 (p<0.01 in all metrics, one-tailed binomial test without ties). It should be noted that both methods identified a large proportion of trash peptides with percentile ranks greater than 20 in the data (∼21% and ∼32% for NetMHCIIpan-4.2 and MixMHC2pred, respectively). This suggests a poor data quality in general, yielding substantially lower performance than observed in our cross-validation. However, the overall performance gain of our method compared to MixMHC2pred suggests that NetMHCIIpan-4.2 is more powerful in the motif deconvolution and identification of DQ ligands.

**Figure 6:**
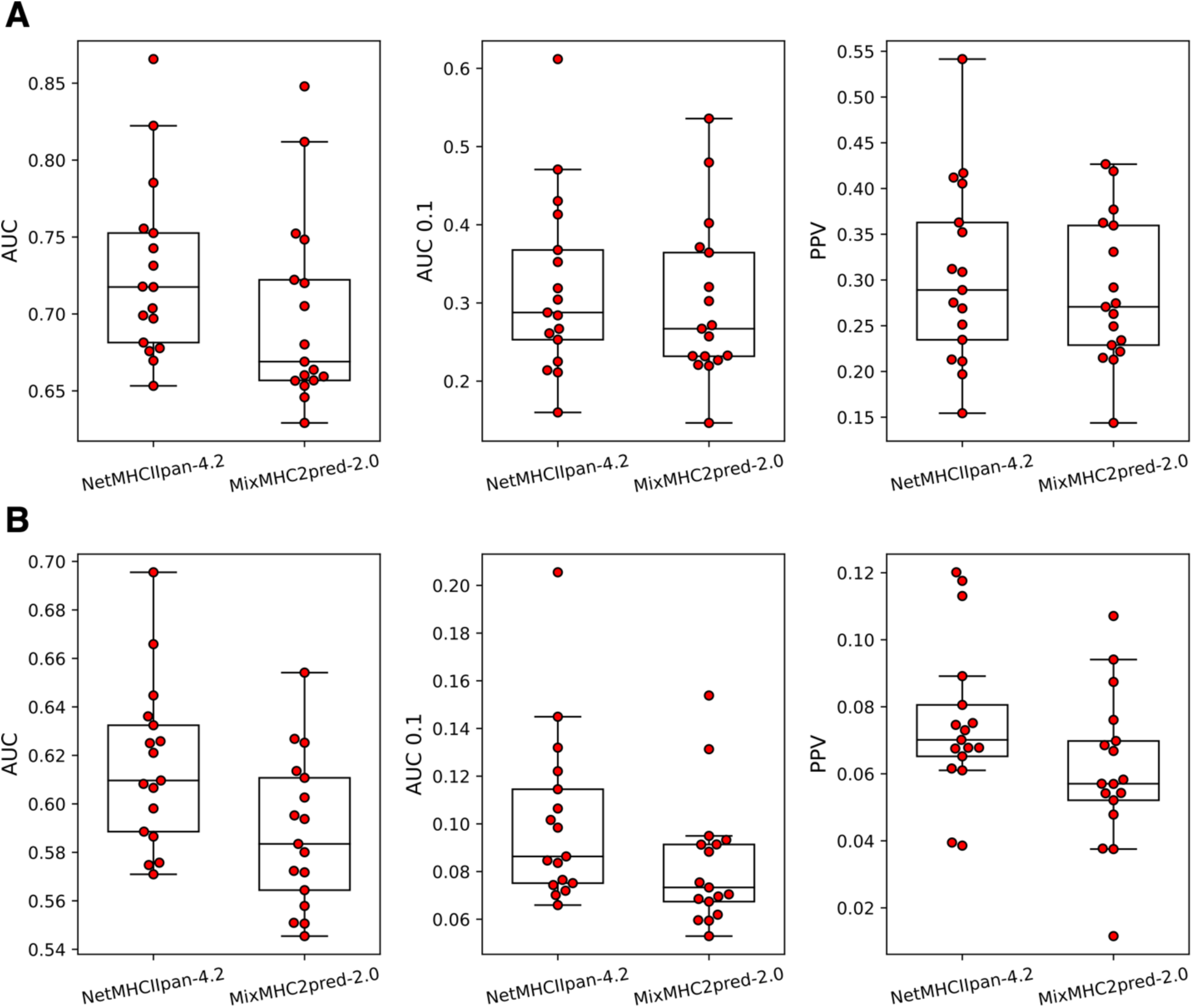
Benchmark against MixMHC2pred-2.0 in terms of AUC, AUC 0.1 and PPV. **A**: Performance per sample calculated on the entire data. **B**: Performance per sample calculated on the union of DQ-annotated peptides between the two methods.

Investigating our method’s motif deconvolution on the DQ-heterozygous samples, we observed that the trans-only molecules once again had limited to no contribution (see supplementary figure 12A). In terms of observed cis variants found in the DQ-SA or DQ-MA training data (cis-SA and cis-MA, respectively), the cis-SA molecules had the largest contribution as expected, with cis-MA having significantly larger contribution than the trans-only variants (p=0.0002, two-sample t-test). Similar results were found when taking into account cis-MA molecules with the same pseudo-sequence as a cis-SA molecule (supplementary figure 12B). This result contrasts with what was observed for MixMHC2pred, where close to an equal contribution was observed across the different molecule classes. Furthermore, when examining the motif deconvolution of the DQ-heterozygous samples without trans-only molecules, our method was able to identify four distinct motifs in each sample (see supplementary figure 12C).

## Discussion

In this work, we have demonstrated how rational data generation combined with refined immunoinformatics data mining can boost the performance of HLA class II antigen presentation predictions and move towards closing the performance gap between HLA-DR and HLA-DQ.

We generated high quality MS-immunopeptidomics data from a series of 16 HLA-DQ homozygous cell lines covering a total of 14 frequent HLA-DQ molecules in different populations worldwide. Using an in-house HLA-DQ specific antibody enabled identification of MS-immunopeptidomics datasets of an, in a DQ context, unprecedented volume with an average of 2,600 unique peptides identified in each cell line. Integrating this large volume of data with earlier data from the development of NetMHCIIpan-4.1 allowed us to boost the HLA-DQ antigen presentation predictive performance to a level comparable to that of HLA-DR. Investigating the accuracy of the motif deconvolution of the two methods trained with and without the novel data demonstrated an overall improved motif consistency across all HLA molecules. This observation demonstrates how integration of the novel HLA-DQ data results in an overall improved HLA-restriction assignment of the individual MS-HLA-peptides leading to more accurate motif characterizations across all three HLA class II loci. The main source of this improvement was demonstrated to be an increased volume of peptide assignment to HLA-DQ molecules during the motif deconvolution. This resulted in improved motif accuracy for both HLA-DQ imposed by the larger volume of peptides, and non HLA-DQ molecules by the removal of peptides mis-assigned as DQ restricted by the model not including the novel DQ data.

Next, moving into the issue of cis versus trans-only HLA-DQ α- and β-chain combinations, we demonstrated that in contrast to the method without the novel data, the model trained including the novel data performed the DQ motif deconvolution almost solely using known HLA-DQ cis-variants. One particular example here was the HLA-DQ molecule DQA1*01:01-DQB1*03:01, which was assigned a large number of peptides in the model trained without the novel data. However, when including the novel data, the peptide assignment to this molecule was almost completely depleted. This result combined with the overall increased HLA-DQ peptide volume and motif consistency of the model trained including the novel data, strongly suggests that trans-only HLA-DQ α and β combinations are non-functional and have minimal or no contribution to the total HLA-DQ immunopeptidome. This finding is striking since the definition of cis and trans-only dimerization defined here precisely follows the rules proposed earlier for forming stable/unstable HLA-DQ heterodimers. Specifically, the rules indicate that structural constraints do not favor dimerization of DQA1*01 with DQB1*02, 03, and 04 alleles, resulting in their inefficient assembly, lack of stability and surface expression and therefore loss of function^12,14^. These results thus demonstrate how such rules can be learned directly from MS-immunopeptidome data using tailored data mining methods and rationally defined data sets, suggesting that similar types of analysis should be extended to HLA-DP to further our understanding of cis versus trans α- and β-chain pairing.

Note, that the definition of cis and trans-only HLA-DQ α- and β-chain combinations applied in this work is contingent on the current haplotype data available and the assumption that all observed haplotype α and β combinations can pair and form cis-variants, and all other combinations not observed as such cis-variants are trans-only. The current data defining these categories are limited in volume, and larger sample sizes are required for more accurate analyses particularly for the more heterogeneous groups and low frequency haplotypes^13^.

Lastly, we demonstrated how the coverage of HLA-DQ molecules was largely increased by the models trained with the novel data and illustrated this by constructing an HLA-DQ tree showing coverage of all branches. This suggests that the current model covers all HLA-DQ binding specificities (considering that *trans-only* HLA-DQ molecules have limited to no contribution to the overall HLA-DQ immunopeptidome).

Overall, this work has demonstrated how careful data generation using a DQ-specific antibody and affinity purification combined with refined data mining and motif deconvolution can be applied towards closing the performance gap in peptide binding prediction between HLA-DR and HLA-DQ. Despite the large performance gain demonstrated here, the accuracy for HLA-DQ remains below what is observed for DR. We demonstrate that this to a very large degree can be attributed to the generally lower quantity and quality of ligands obtained in earlier DQ immunoprecipitation studies where most often DQ (and DP) data have been obtained using a pan-HLA class II antibody (after first depleting for HLA-DR^29^). Focusing solely on the novel data generated in this study, we find that both the quantity and quality of the obtained DQ ligands are on par with what is found for HLA-DR, resulting in predictive performance for the associated dataset being equal between the two. This result has large impacts and suggests that modeling DQ is a task of equal complexity to that of HLA-DR, and that the current lower performance of DQ compared to DR is driven by low quantity and quality of data; a situation that can be resolved by generation of high quality and volume data as outlined in this study.

In conclusion, other than demonstrating an overall improved predictive performance and coverage of HLA-DQ molecules, a key result of our work is an improved understanding of the relative contribution of cis versus trans-only paired molecules to the total HLA-DQ immunopeptidome demonstrating a very limited role of the latter in complementing the specificity space. We believe these findings will provide a foundation for further research defining the molecular role of HLA-DQ in the onset of cellular immunity within autoimmune and infectious diseases.

## Acknowledgements

We would like to sincerely thank Dr. Rico Buchli (Pure Protein, LLC) for providing the SPVL3 affinity columns for this study. We also thank Steven Cate (University of Oklahoma Health Sciences Center) and Sean Osborn (Pure MHC, LLC) for HLA typing of the BLCLs and very helpful discussions.

## Author contributions

SK and MN designed the study. The experimental data used in the study was generated by SK, with contribution from HY and WH. JN and MN generated the computational results and figures. CB, LG and BP contributed towards methodology and provided scientific feedback. The manuscript was written by JN, SK and MN, with contributions from all authors. All authors have read and approved the final version of the paper.

## Competing interests

SK is an employee at Pure MHC, LLC. The remaining authors declare no competing interests.

## Data availability

The novel immunopeptidomics data generated for this study is available in supplementary table 2 in the manuscript (provided in saghar_peptide_data.xlsx).

## Funding information

Research reported in this publication was supported by the National Institute of Allergy and Infectious Diseases (NIAID), under award number 75N93019C00001.

## Supplementary figures

**Supplementary figure 1:**
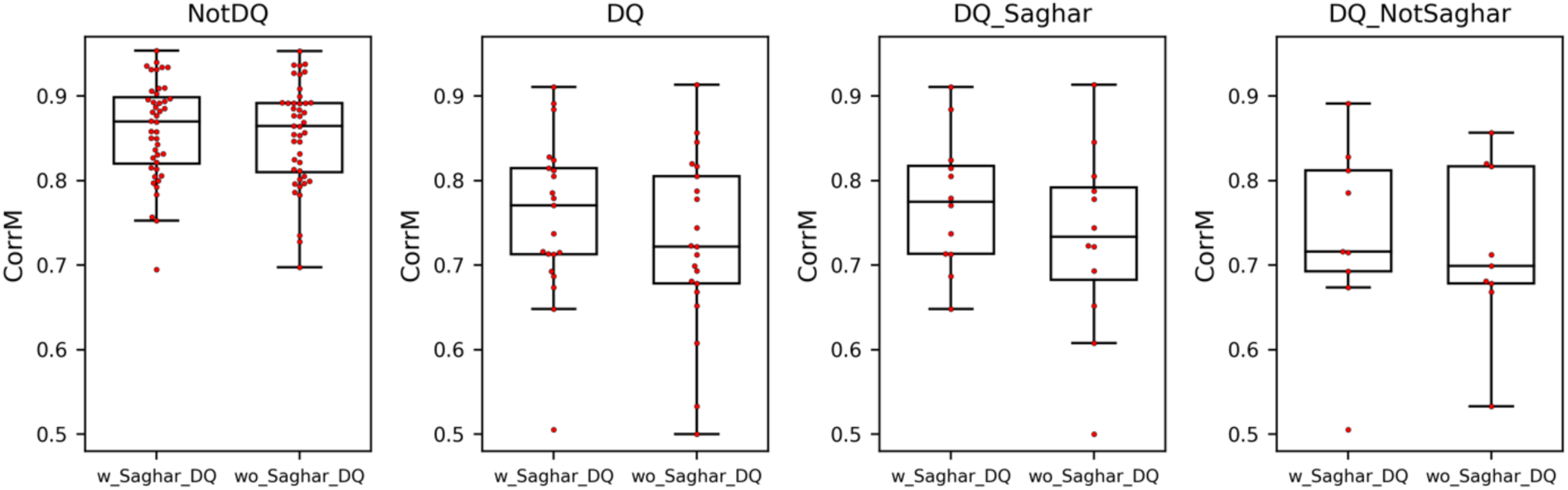
Mean consistency values for HLA molecule deconvolutions shared between multiple data sets for the two models. Each point reflects an HLA class II molecule, and values shown are the mean over the different data set comparisons. The different subgroups of HLA molecules are defined in the main text and the Figure 2 caption.

**Supplementary figure 2:**
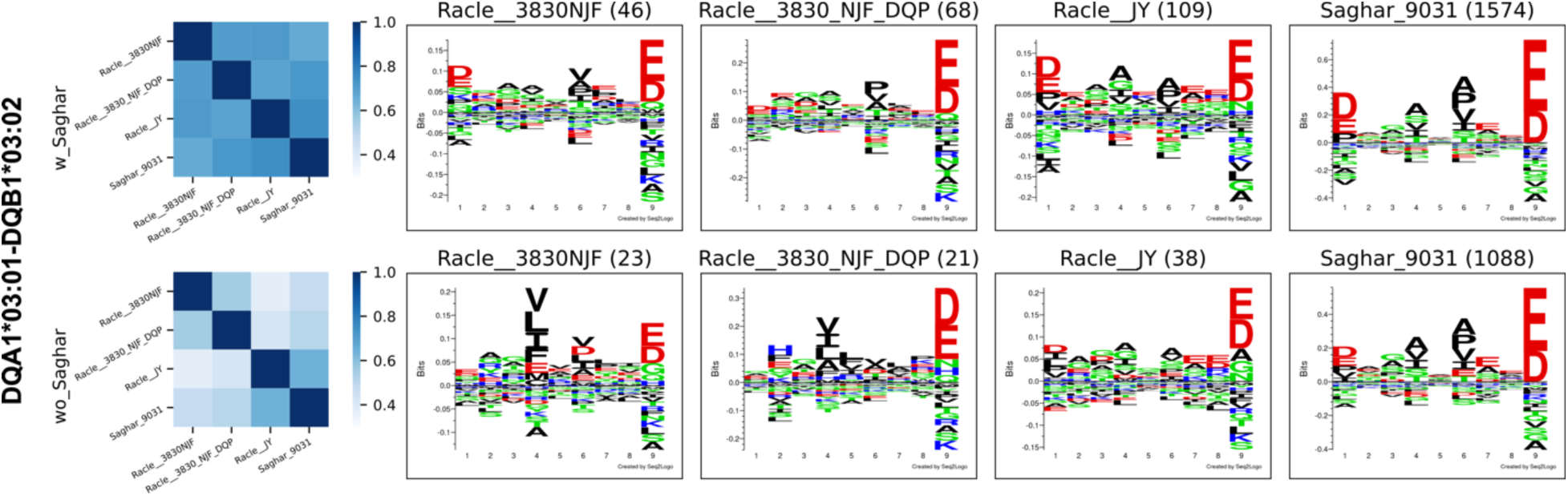
Consistency analysis for HLA-DQA1*03:01-DQB1*03:02. Here, only a subset of the cell lines used in the consistency analysis are included. Left panel: Correlation matrices showing the consistency values per cell line pair. Right panel: Sequence logos constructed from the peptide sets used in the consistency correlation analysis, with a minimum threshold of 20 peptides required for a given logo. The rows correspond to the methods including (w_Saghar) and not including (wo_Saghar) the novel data. The peptide count for each logo is shown in parenthesis in the logo title.

**Supplementary figure 3:**
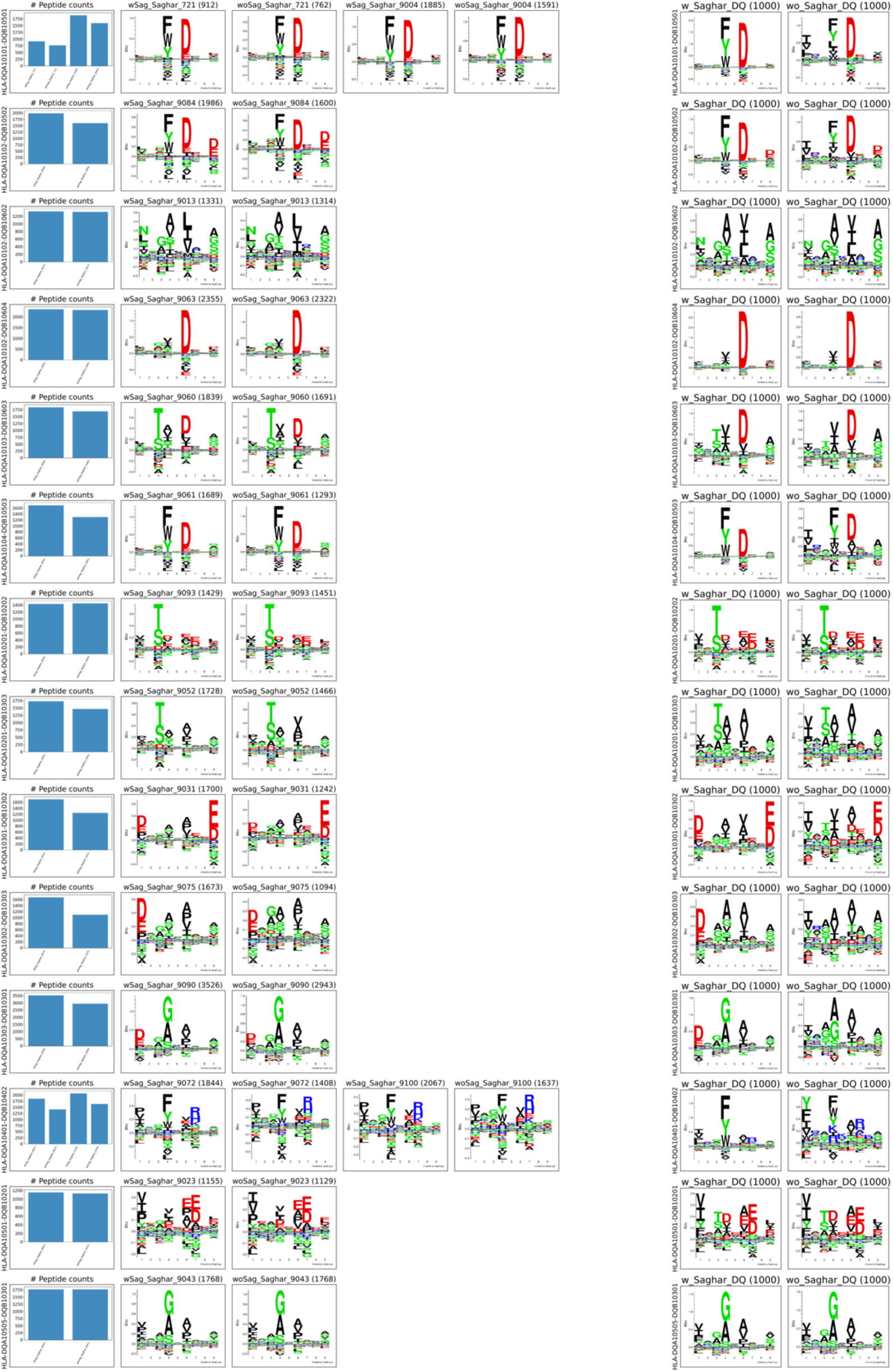
Identified binding motifs for the 14 DQ molecules from the novel data in the two methods. Left panel: Motif deconvolution of the novel datasets in the models trained with (wSag) and without (woSag) the novel data (the two rows with 4 logos correspond to the two HLA-DQ molecules shared between two cell lines). Right panel: Motifs predicted for the HLA-DQ molecules by the two models. Predicted motifs are generated from the top 1% of 100,000 random natural 13-17 mer peptides.

**Supplementary figure 4:**
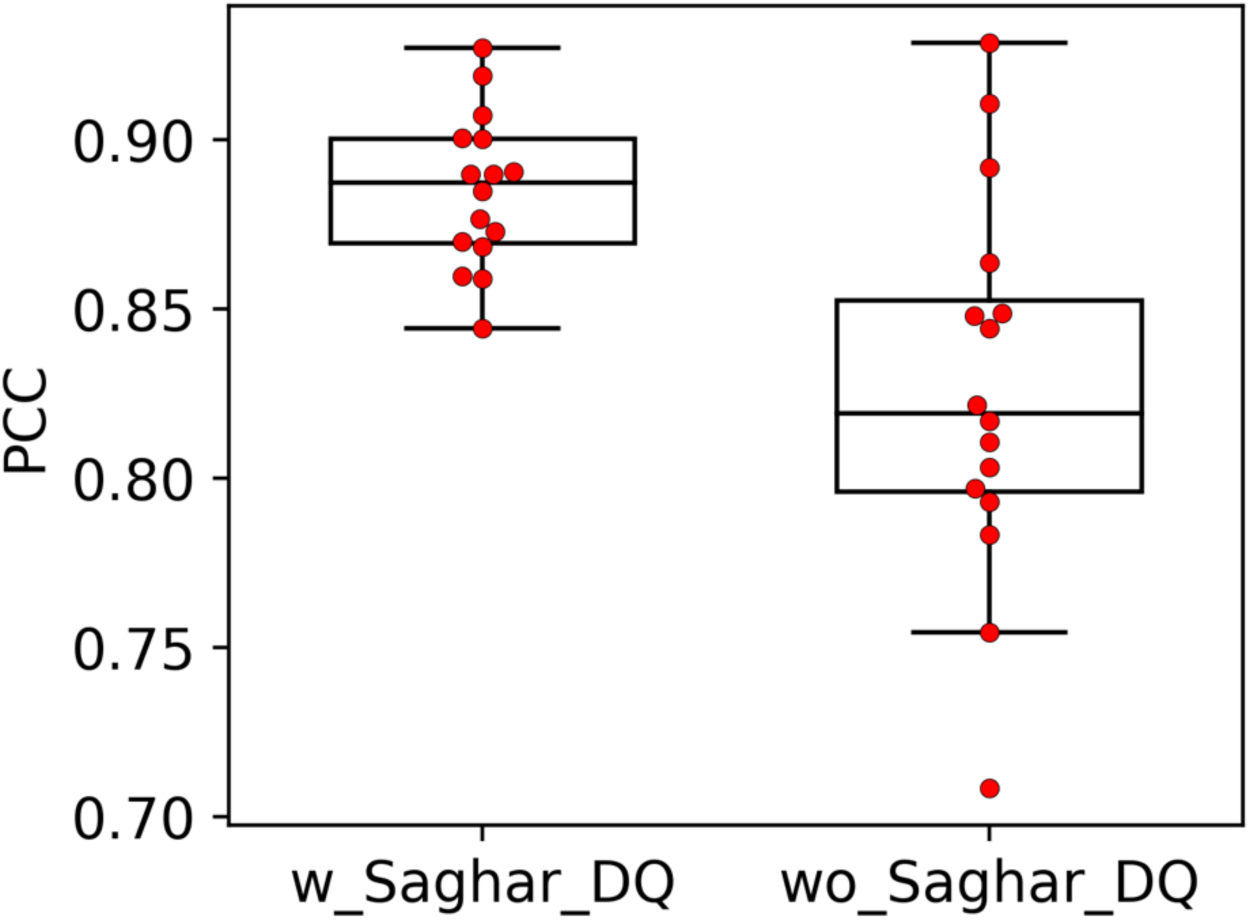
Correlations between deconvoluted and predicted sequence logos in the methods with (w_Saghar) and without (wo_Saghar) the novel data. For each cell line in the novel data, the PCC value between the DQ motif deconvolution PSSM and predicted PSSM for the corresponding molecule was calculated for each of the methods with and without the novel data.

**Supplementary figure 5:**
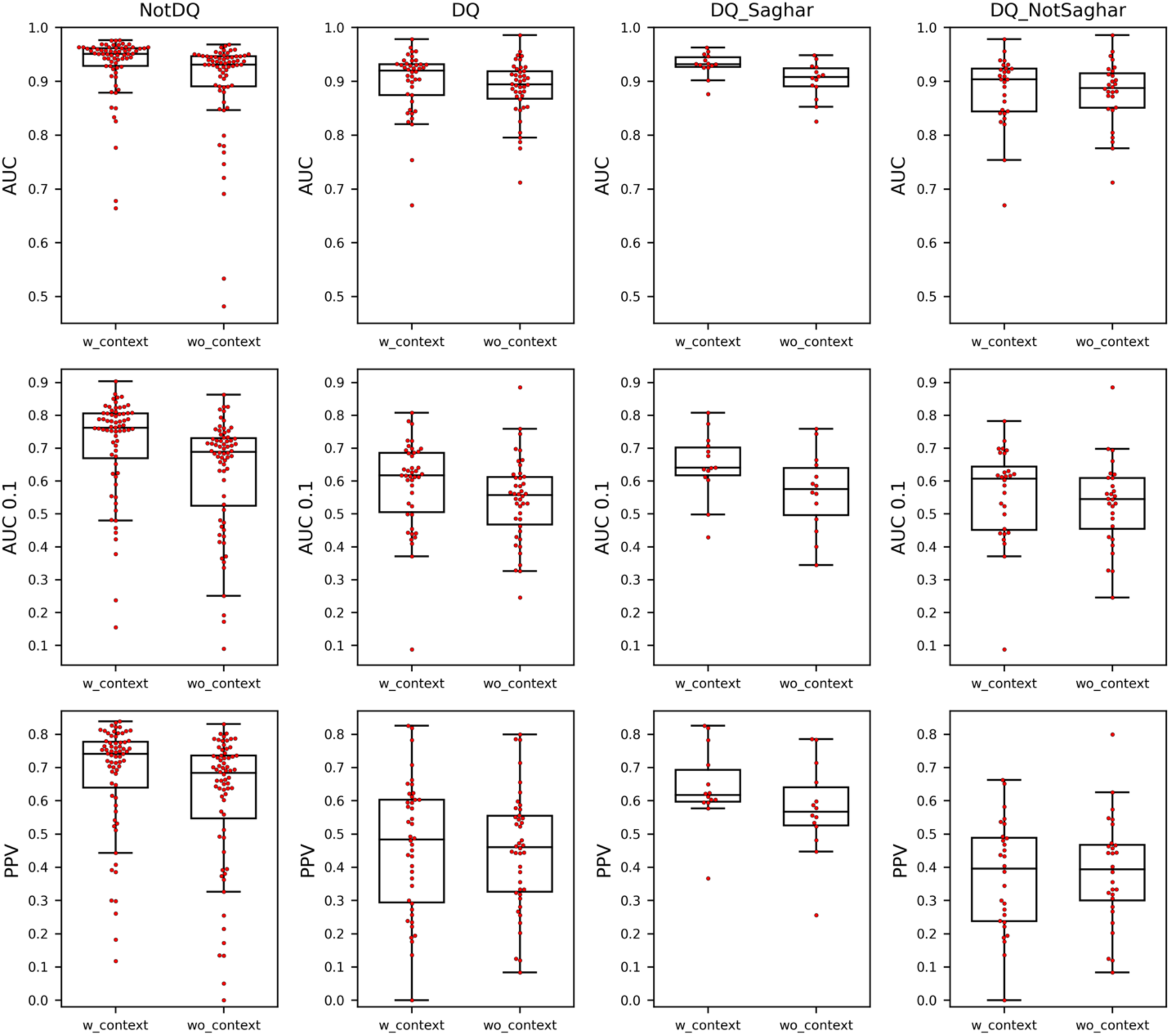
Performance comparison of models trained with the novel DQ data, with (w_context) and without (wo_context) context encoding. The different subgroups of HLA molecules are defined in the main text and the Figure 2 caption.

**Supplementary figure 6:**
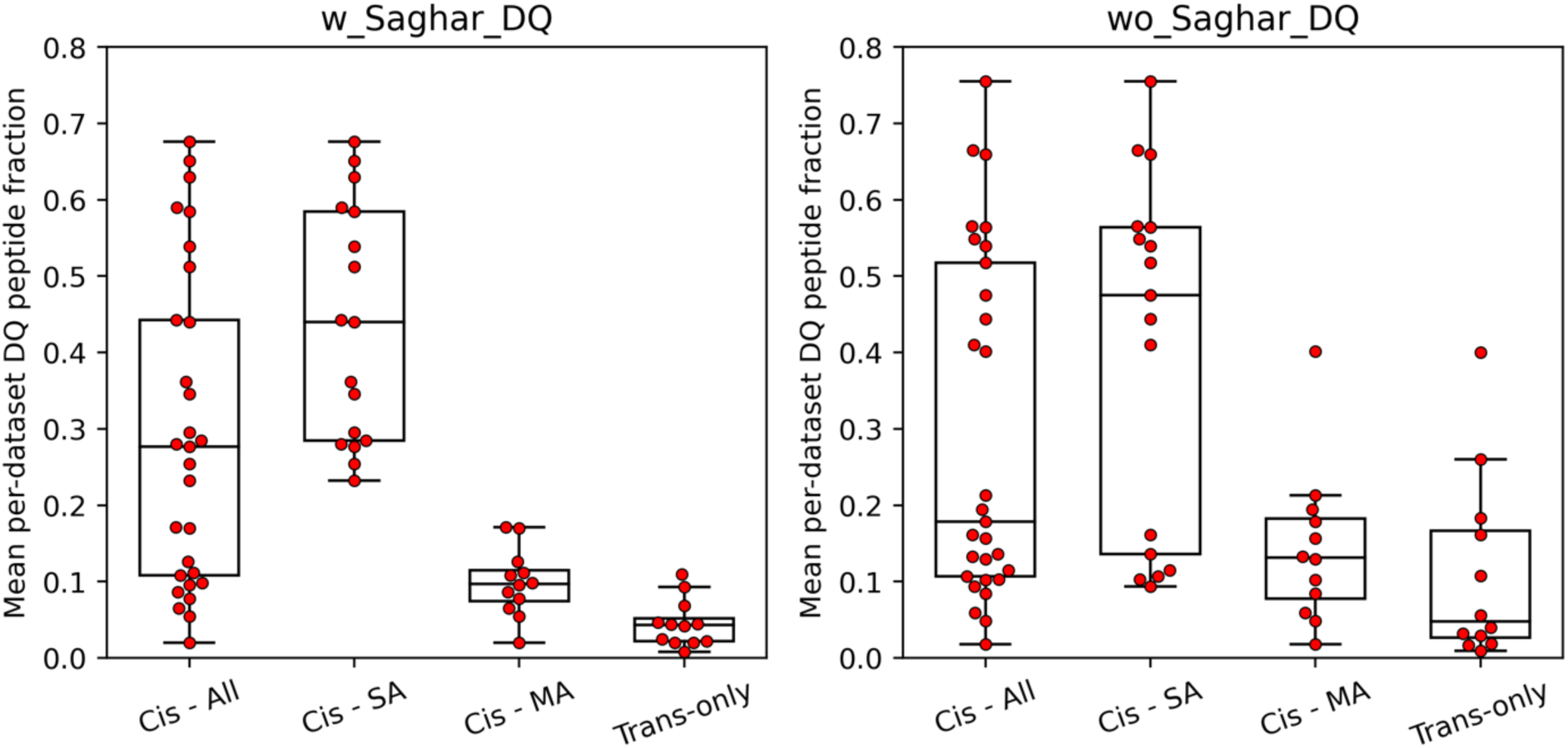
Peptide-count contribution of cis and trans-only molecules in the methods with (w_Saghar_DQ) and without (wo_Saghar_DQ) the novel data, taking into account pseudo-sequence overlap. Each point shows the mean per-dataset peptide fraction for a given DQ molecule. The cis molecules are shown in three categories, namely all cis molecules (Cis - All), cis molecules found in the DQ-SA training data or with the same pseudosequence as a DQ-SA molecule (Cis - SA), and cis molecules only found in the DQ-MA training data and with no pseudosequence overlap to cis-SA molecules (Cis - MA). Here, a significant difference was found between cis-MA and trans-only in the model with the novel data (p=0.002, two-sample t-test).

**Supplementary figure 7:**
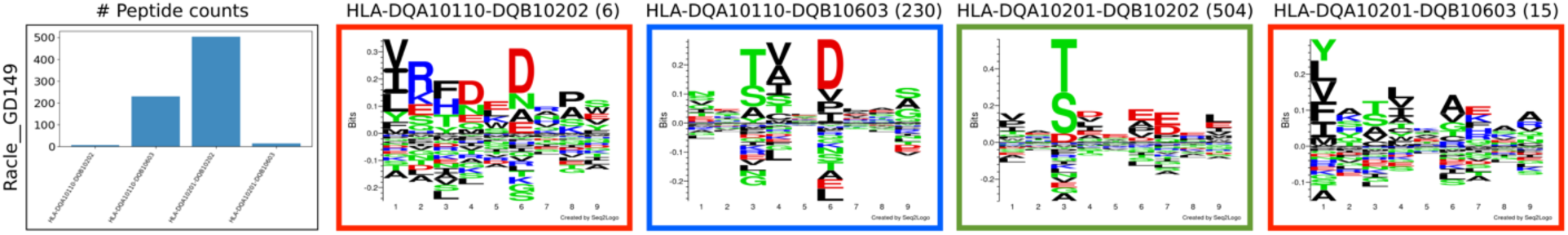
DQ motif deconvolution of the Racle GD149 dataset. The logo frame colors indicate three types of DQ molecules, namely cis-molecules present in the DQ-SA training data (green), cis molecules present in the DQ-MA training data (blue), and trans-only molecules (red).

**Supplementary figure 8:**
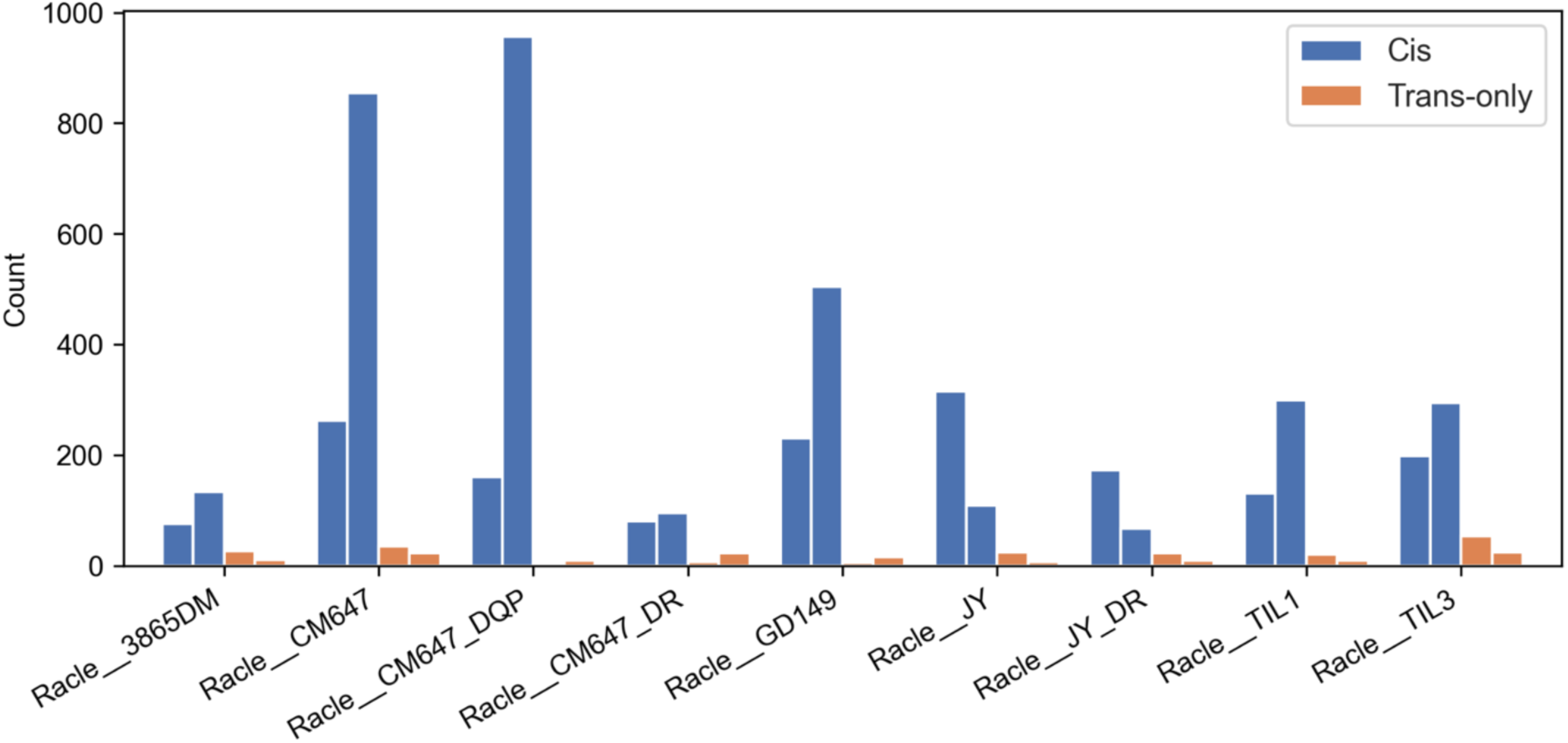
DQ ligand count distributions in DQ-heterozygous cell lines with both cis and trans-only molecules. Only cell lines with at least 100 DQ-annotated peptides excluding trash are included. The counts are shown for the individual molecules, colored by category (cis: blue and trans-only: orange). If we assume that the relative expression from each chromosome is linked to the observed peptide counts for the cis variants, then the trans-only variants (if they were functional) should in principle have peptide counts corresponding to at least the cis variant with the lowest count. However, this is not what we observe, as the cis variant with the lowest count always has a much larger peptide count than both trans-only variants.

**Supplementary figure 9:**
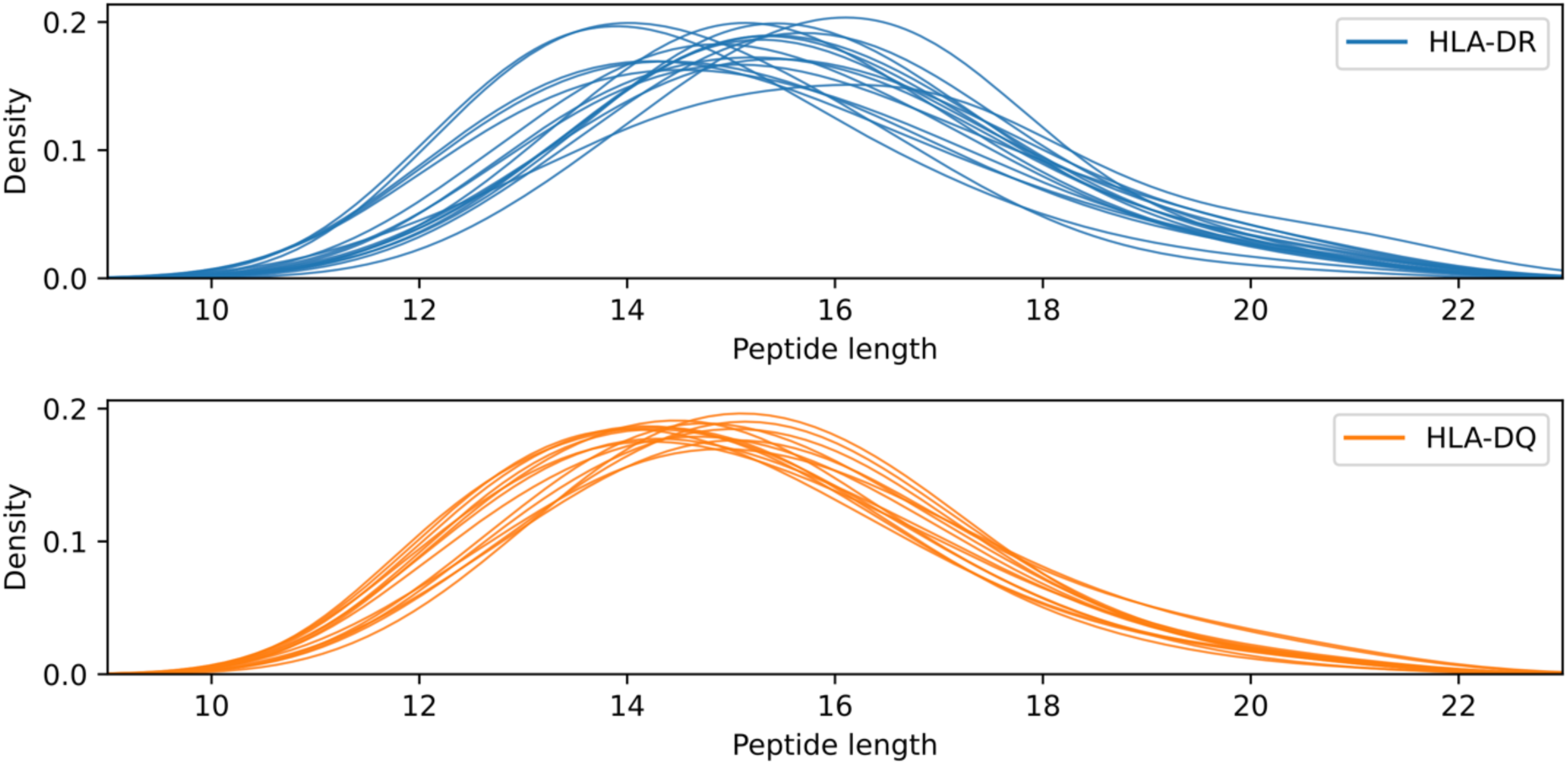
Kernel density plots of length distributions of peptides extracted from the cell lines used in the novel data. Each curve corresponds to the peptide length distribution for an HLA-DR (top) or HLA-DQ (bottom) molecule. The DR distributions are based on the immunopeptidomics data from supplementary table 4 in Kaabinejadian et al. 2022, which were run through the motif deconvolution method developed in this manuscript.

**Supplementary figure 10:**
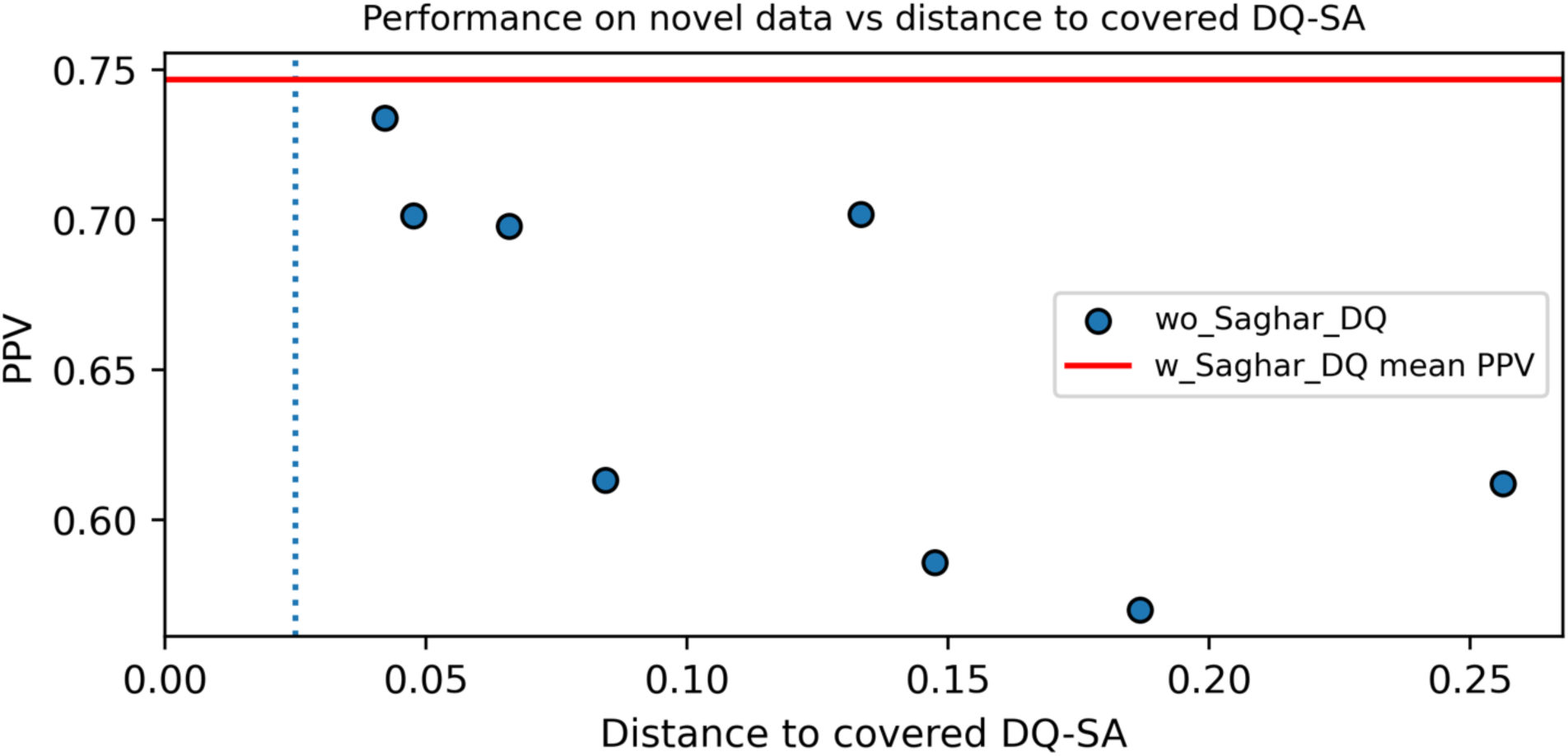
Performance vs distance analysis for the model trained without the novel data (wo_Saghar_DQ). Each point shows the PPV performance evaluated on the novel data for a DQ molecule not present in the DQ-SA training data of the wo_Saghar_DQ method, as a function of the distance to the training data DQ-SA molecules. The red line shows the mean performance of the model trained with the novel data (w_Saghar_DQ) on the same set of DQ molecules, evaluated on the novel data.

**Supplementary figure 11:**
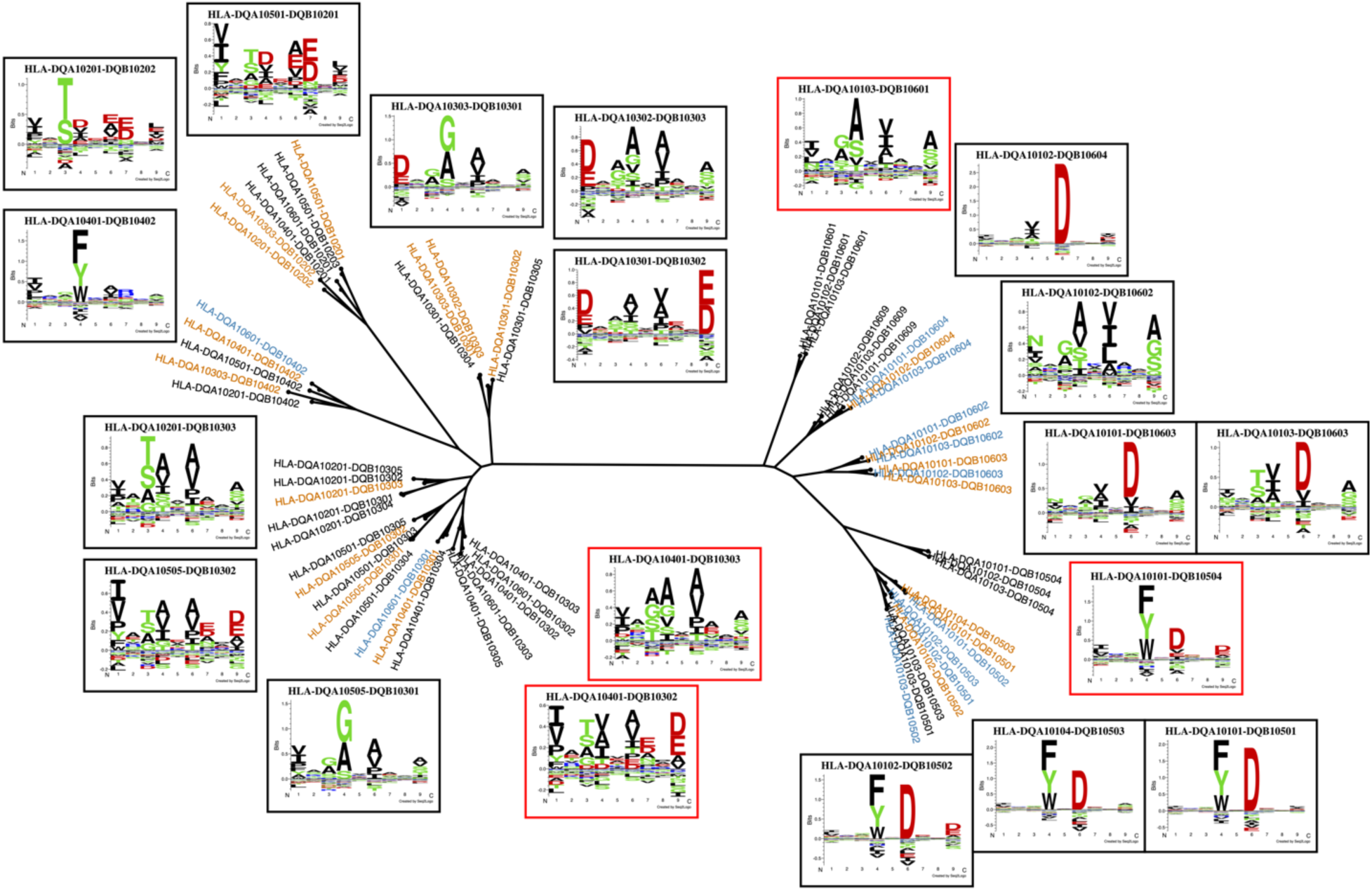
Sequence-based clustering of DQ molecules. The tree is based on 61 DQ molecules including the 14 molecules described by the novel data. Orange molecules are covered by the method including the novel data with at least 100 peptides, and blue molecules are within a distance 0.025 of an orange molecule. Black molecules are non-covered (i.e. have peptide count less than 100 and have distance greater than 0.025 to an orange molecule). Logos in black frames correspond to orange molecules. Logos in red frames correspond to molecules from branches with clusters of non-covered (black) molecules. The phylogenetic tree was constructed from the DQ pseudo-sequences using ClustalW. Logos were constructed from the top 1% of 100,000 random 13-17 mer peptides.

**Supplementary figure 12A:**
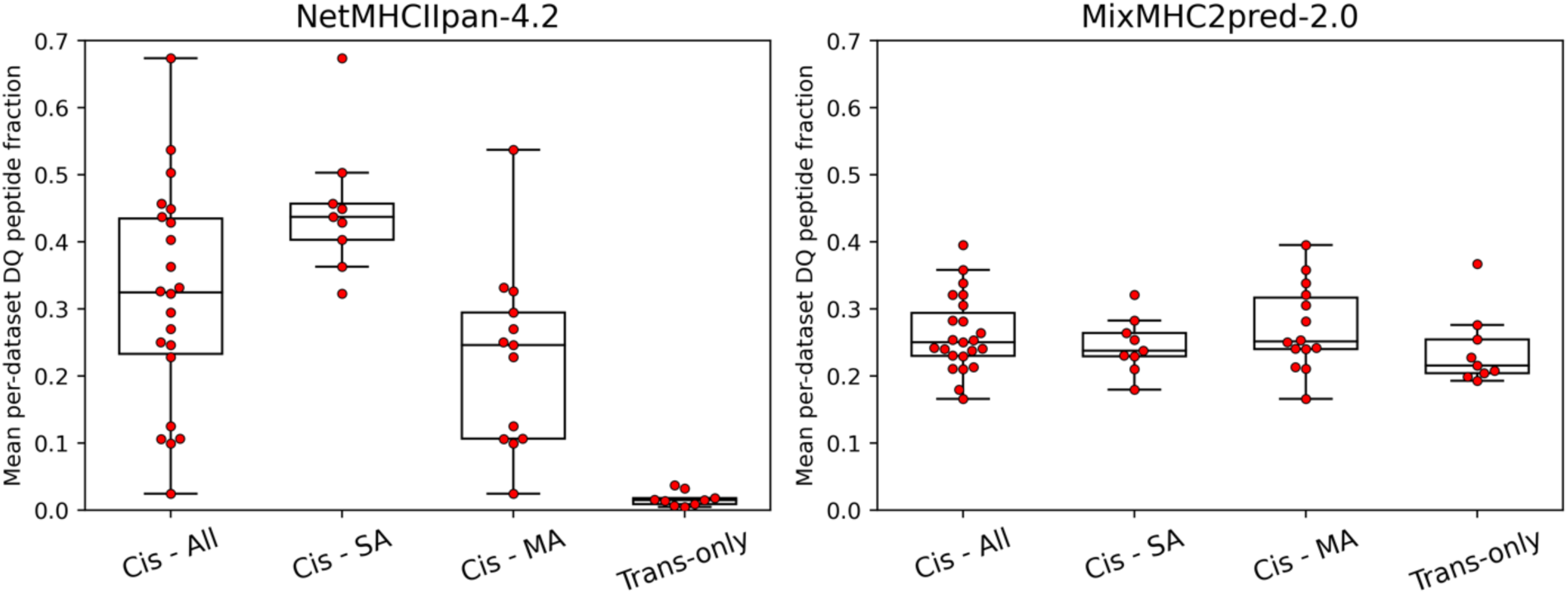
Peptide-count contribution of cis and trans-only molecules predicted by NetMHCIIpan-4.2 and MixMHC2pred-2.0 on DQ-heteryzygous data from Marcu et al. 2021. Each point shows the mean per-dataset peptide fraction for a given DQ molecule. The cis molecules are shown in three categories, namely all cis molecules (Cis - All), cis molecules found in the DQ-SA training data (Cis - SA), and cis molecules only found in the DQ-MA training data (Cis - MA).

**Supplementary figure 12B:**
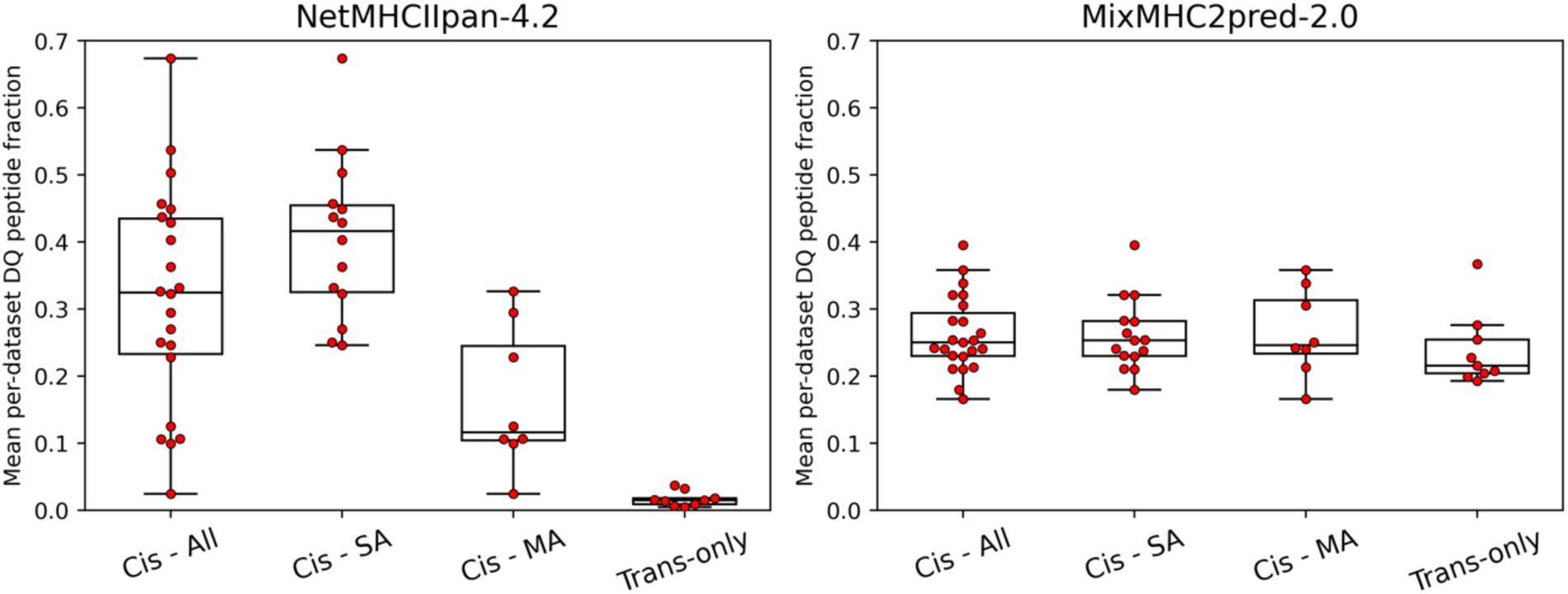
Peptide-count contribution of cis and trans-only molecules predicted by NetMHCIIpan-4.2 and MixMHC2pred-2.0 on DQ-heteryzygous data from Marcu et al. 2021, taking into account pseudo-sequence overlap. Each point shows the mean per-dataset peptide fraction for a given DQ molecule. The cis molecules are shown in three categories, namely all cis molecules (Cis - All), cis molecules found in the DQ-SA training data or with the same pseudosequence as a DQ-SA molecule (Cis - SA), and cis molecules only found in the DQ-MA training data and with no pseudosequence overlap to cis-SA molecules (Cis - MA). Here, a significant difference was found between cis-MA and trans-only in NetMHCIIpan-4.2 (p=0.001, two-sample t-test).

**Supplementary figure 12C:**
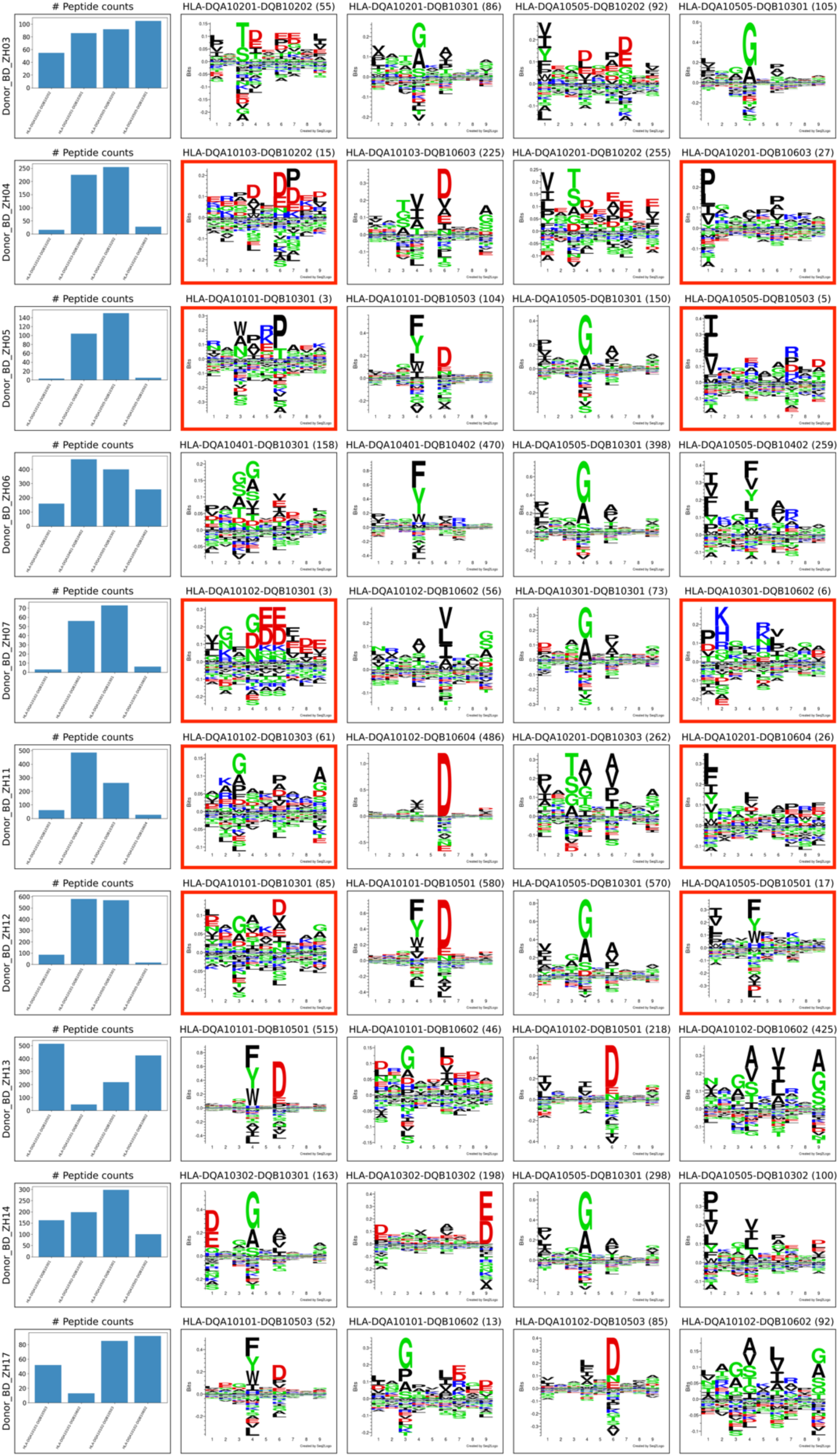
DQ motif deconvolution for DQ-heterozygous datasets in the benchmark data from Marcu et al. 2021. Each row corresponds to a donor sample. Only peptides with percentile rank less than 5 were included in the logo plots. Trans-only molecules are highlighted in red frames.

## Supplementary tables

**Supplementary table 1:** Overview of HLA types per EL dataset used to train our method. The novel data generated for this study are highlighted in bold. For two of the datasets from Abelin et al. 2019 (Abelin HLA_DR_SILAC_Donor2_DC_UV_16hr and Abelin HLA_DR_SILAC_Donor2_DC_UV_24hr), the original HLA typing was listed to have only two DQ molecules. However, we believe this to be an erroneous HLA typing, as the cell lines should express four different DQ molecules. As such, the HLA typing for these datasets was extended to include all four DQ heterodimers.

| Sample | HLA type |
| --- | --- |
| Saghar_721 | DRB1*01:01,DQA1*01:01-DQB1*05:01,DPA1*01:03-DPB1*02:01 |
| Saghar_9004 | DRB1*01:01,DQA1*01:01-DQB1*05:01,DPA1*01:03-DPB1*04:01 |
| Saghar_9013 | DRB1*15:01,DRB5*01:01,DQA1*01:02-DQB1*06:02,DPA1*01:03-DPB1*04:02 |
| Saghar_9023 | DRB1*03:01,DRB3*01:01,DQA1*05:01-DQB1*02:01,DPA1*02:01-DPB1*01:01 |
| Saghar_9031 | DRB1*04:01,DRB4*01:03,DQA1*03:01-DQB1*03:02,DPA1*01:03-DPB1*04:01 |
| Saghar_9043 | DRB1*11:01,DRB3*02:02,DQA1*05:05-DQB1*03:01,DPA1*02:01-DPB1*10:01 |
| Saghar_9052 | DRB1*07:01,DRB4*01:03N,DQA1*02:01-DQB1*03:03,DPA1*01:03-DPB1*04:01 |
| Saghar_9060 | DRB1*13:01,DRB3*02:02,DQA1*01:03-DQB1*06:03,DPA1*02:02-DPB1*19:01 |
| Saghar_9061 | DRB1*14:01,DRB3*02:02,DQA1*01:04-DQB1*05:03,DPA1*01:03-DPB1*04:01 |
| Saghar_9063 | DRB1*13:02,DRB3*03:01,DQA1*01:02-DQB1*06:04,DPA1*01:03-DPB1*16:01 |
| Saghar_9072 | DRB1*08:02,DQA1*04:01-DQB1*04:02,DPA1*01:03-DPB1*04:02 |
| Saghar_9075 | DRB1*09:01,DRB4*01:03,DQA1*03:02-DQB1*03:03,DPA1*01:03-DPB1*04:01 |
| Saghar_9084 | DRB1*16:01,DRB5*02:02,DQA1*01:02-DQB1*05:02,DPA1*01:03-DPB1*04:01 |
| Saghar_9090 | DRB1*04:01,DRB4*01:03,DQA1*03:03-DQB1*03:01,DPA1*01:03-DPB1*04:01,DPA1*01:03-DPB1*14:01,DPA1*02:01-DPB1*04:01,DPA1*02:01-DPB1*14:01 |
| Saghar_9093 | DRB1*07:01,DRB4*01:03,DQA1*02:01-DQB1*02:02,DPA1*01:03-DPB1*02:01,DPA1*01:03-DPB1*04:01 |
| Saghar_9100 | DRB1*08:02,DQA1*04:01-DQB1*04:02,DPA1*01:03-DPB1*03:01,DPA1*01:03-DPB1*04:02 |
| Abelin__MAPTAC_DRB1_0403 | DRB1*04:03 |
| Abelin__MAPTAC_DRB1_0803 | DRB1*08:03 |
| Abelin__MAPTAC_DRB5_0101 | DRB5*01:01 |
| Abelin__MAPTAC_DRB3_0202 | DRB3*02:02 |
| Abelin__MAPTAC_DPB1_0601_DPA1_0103_dmPos | DPA1*01:03-DPB1*06:01 |
| Abelin__MAPTAC_DQB1_0604_DQA1_0102_dmPos | DQA1*01:02-DQB1*06:04 |
| Abelin__MAPTAC_HLA_DPB10201_DPA10103 | DPA1*01:03-DPB1*02:01 |
| Abelin__MAPTAC_HLA_DQB10602_DQA10102 | DQA1*01:02-DQB1*06:02 |
| Abelin__MAPTAC_DRB1_0101 | DRB1*01:01 |
| Abelin__MAPTAC_DRB1_0102 | DRB1*01:02 |
| Abelin__MAPTAC_DRB1_0301 | DRB1*03:01 |
| Abelin__MAPTAC_DRB1_0302 | DRB1*03:02 |
| Abelin__MAPTAC_DRB1_0401 | DRB1*04:01 |
| Abelin__MAPTAC_DRB1_0402 | DRB1*04:02 |
| Abelin__MAPTAC_DRB1_0404 | DRB1*04:04 |
| Abelin__MAPTAC_DRB1_0405 | DRB1*04:05 |
| Abelin__MAPTAC_DRB1_0407 | DRB1*04:07 |
| Abelin__MAPTAC_DRB1_0701 | DRB1*07:01 |
| Abelin__MAPTAC_DRB1_0801 | DRB1*08:01 |
| Abelin__MAPTAC_DRB1_0802 | DRB1*08:02 |
| Abelin__MAPTAC_DRB1_0804 | DRB1*08:04 |
| Abelin__MAPTAC_DRB1_0901 | DRB1*09:01 |
| Abelin__MAPTAC_DRB1_1001 | DRB1*10:01 |
| Abelin__MAPTAC_DRB1_1101 | DRB1*11:01 |
| Abelin__MAPTAC_DRB1_1102 | DRB1*11:02 |
| Abelin__MAPTAC_DRB1_1104 | DRB1*11:04 |
| Abelin__MAPTAC_DRB1_1202 | DRB1*12:02 |
| Abelin__MAPTAC_DRB1_1301 | DRB1*13:01 |
| Abelin__MAPTAC_DRB1_1302 | DRB1*13:02 |
| Abelin__MAPTAC_DRB1_1303 | DRB1*13:03 |
| Abelin__MAPTAC_DRB1_1401 | DRB1*14:01 |
| Abelin__MAPTAC_DRB1_1501 | DRB1*15:01 |
| Abelin__MAPTAC_DRB1_1502 | DRB1*15:02 |
| Abelin__MAPTAC_DRB1_1503 | DRB1*15:03 |
| Abelin__MAPTAC_DRB1_1601 | DRB1*16:01 |
| Abelin__MAPTAC_DRB3_0301 | DRB3*03:01 |
| Abelin__MAPTAC_DRB4_0103 | DRB4*01:03 |
| Abelin__MAPTAC_DRB1_1201 | DRB1*12:01 |
| Abelin__MAPTAC_DRB3_0101 | DRB3*01:01 |
| Abelin__HLA_DR_A375 | DRB1*07:01,DRB4*01:01,DRB1*04:05 |
| Abelin__HLA_DR_Lung | DRB1*01:01,DRB1*03:01,DRB3*01:01,DPA1*01:03-DPB1*02:01,DPA1*02:01-DPB1*02:01,DPA1*01:03-DPB1*10:01,DPA1*02:01-DPB1*10:01,DQA1*01:01-DQB1*02:01,DQA1*01:01-DQB1*05:01,DQA1*05:01-DQB1*02:01,DQA1*05:01-DQB1*05:01 |
| Abelin__HLA_DR_PBMCD_HDSC | DRB1*03:01,DRB1*11:01,DRB3*01:01,DRB3*02:02 |
| Abelin__HLA_DR_PBMCD_RG1095 | DRB1*03:01,DRB1*11:01,DRB3*01:01,DRB3*02:02 |
| Abelin__HLA_DR_PBMCD_RG1104 | DRB1*01:01,DRB1*11:01,DRB3*02:02 |
| Abelin__HLA_DR_PBMCD_RG1248 | DRB1*03:01,DRB3*01:01 |
| Abelin__HLA_DR_SILAC_Donor1_10minLysate | DRB1*07:01,DRB4*01:01 |
| Abelin__HLA_DR_SILAC_Donor1_5hrLysate | DRB1*07:01,DRB4*01:01 |
| Abelin__HLA_DR_SILAC_Donor1_DConly | DRB1*07:01,DRB4*01:01 |
| Abelin__HLA_DR_SILAC_Donor1_UVovernight | DRB1*07:01,DRB4*01:01 |
| Abelin__HLA_DR_SILAC_Donor2_DC_UV_16hr | DRB1*04:01,DRB4*01:03,DRB1*15:03,DRB5*01:01,DQA1*03:01-DQB1*03:02,DQA1*01:02-DQB1*06:02,DQA1*01:02-DQB1*03:02,DQA1*03:01-DQB1*06:02,DPA1*01:03-DPB1*02:01,DPA1*01:03-DPB1*04:01 |
| Abelin__HLA_DR_SILAC_Donor2_DC_UV_24hr | DRB1*04:01,DRB4*01:03,DRB1*15:03,DRB5*01:01,DQA1*03:01-DQB1*03:02,DQA1*01:02-DQB1*06:02,DQA1*01:02-DQB1*03:02,DQA1*03:01-DQB1*06:02,DPA1*01:03-DPB1*02:01,DPA1*01:03-DPB1*04:01 |
| Abelin__HLA_DR_Spleen | DRB1*01:01,DRB1*03:01,DRB3*01:01,DPA1*01:03-DPB1*02:01,DPA1*02:01-DPB1*02:01,DPA1*01:03-DPB1*10:01,DPA1*02:01-DPB1*10:01,DQA1*01:01-DQB1*02:01,DQA1*01:01-DQB1*05:01,DQA1*05:01-DQB1*02:01,DQA1*05:01-DQB1*05:01 |
| Benito__T2_DR3 | DRB1*03:01,DRB3*01:01 |
| Bergseng__9037_SWEIG | DRB1*11:01,DRB3*02:02,DQA1*05:05-DQB1*03:01,DPA1*01:03-DPB1*04:02 |
| Bergseng__9047_PLH | DRB1*07:01,DRB4*01:01,DQA1*02:01-DQB1*02:02,DPA1*01:03-DPB1*15:01 |
| Bergseng__9050_MOU | DRB1*07:01,DRB4*01:01,DQA1*02:01-DQB1*02:02,DPA1*01:03-DPB1*02:01 |
| Bergseng__9051_PITOUT | DRB1*07:01,DRB4*01:01,DQA1*02:01-DQB1*02:02,DPA1*01:03-DPB1*04:01 |
| Bergseng__9064_AMALA | DRB1*14:02,DRB3*01:01,DQA1*05:01-DQB1*03:01,DQA1*05:03-DQB1*03:01,DPA1*01:03-DPB1*04:02,DPA1*01:03-DPB1*94:01 |
| Bergseng__9087_STEINLIN | DRB1*03:01,DRB3*01:01,DQA1*05:01-DQB1*02:01,DPA1*01:03-DPB1*03:01,DPA1*01:03-DPB1*04:01 |
| Bergseng__9088_PFO4015 | DRB1*03:01,DRB3*01:01,DQA1*05:01-DQB1*02:01,DPA1*01:03-DPB1*01:01,DPA1*02:01-DPB1*04:01,DPA1*01:03-DPB1*04:01,DPA1*02:01-DPB1*01:01 |
| Bergseng__9089_BOB | DRB1*11:04,DRB3*02:02,DQA1*05:05-DQB1*03:01,DPA1*01:03-DPB1*04:02 |
| Clement__DRB1_0101 | DRB1*01:01 |
| Heyder__Bcell | DRB1*04:01,DRB4*01:01 |
| Heyder__PAT1 | DRB1*04:01,DRB1*07:01,DRB4*01:01 |
| Heyder__PAT7 | DRB1*03:01,DRB1*15:01,DRB3*01:01,DRB5*01:01 |
| Karuna__H_2_IAb | H-2-IAb |
| Khoda__Jeko | DRB1*04:03,DRB1*07:01,DRB4*01:01 |
| Khoda__L128 | DRB1*07:01,DRB1*11:01,DRB4*01:01,DRB3*02:02 |
| Khoda__MCL001 | DRB1*04:02,DRB1*07:01,DRB4*01:01 |
| Khoda__MCL005 | DRB1*04:01,DRB1*11:04,DRB4*01:01,DRB3*02:02 |
| Khoda__MCL008 | DRB1*04:01,DRB1*13:01,DRB4*01:01,DRB3*01:01 |
| Khoda__MCL022 | DRB1*01:03,DRB1*04:01,DRB4*01:01 |
| Khoda__MCL030 | DRB1*07:01,DRB1*13:03,DRB4*01:01 |
| Khoda__MCL034 | DRB1*03:01,DRB1*04:01,DRB3*01:01,DRB4*01:01 |
| Khoda__MCL037 | DRB1*11:01,DRB1*14:54,DRB3*02:02 |
| Khoda__MCL041 | DRB1*04:01,DRB1*13:01,DRB4*01:01,DRB3*01:01 |
| Khoda__MCL043 | DRB1*07:01,DRB1*11:01,DRB4*01:01,DRB3*02:02 |
| Khoda__MCL049 | DRB1*01:01,DRB1*07:01,DRB4*01:01 |
| Khoda__MCL052 | DRB1*01:01,DRB1*13:01,DRB3*01:01 |
| Khoda__MCLX001 | DRB1*01:01,DRB1*07:01,DRB4*01:01 |
| Khoda__MCLX002 | DRB1*07:01,DRB1*13:01,DRB4*01:01,DRB3*01:01 |
| Nanaware__WT_1 | DRB1*01:01,DQA1*01:01-DQB1*05:01,DPA1*01:03-DPB1*04:01 |
| Nanaware__WT_2 | DRB1*01:01,DQA1*01:01-DQB1*05:01,DPA1*01:03-DPB1*04:01 |
| Nanaware__WT_H2 | H-2-IAb |
| Nelde__UPN1 | DRB1*08:03,DRB1*13:01,DRB3*01:01 |
| Nelde__UPN2 | DRB1*08:01,DRB1*13:02,DRB3*03:01 |
| Nelde__UPN3 | DRB1*07:01,DRB1*08:01,DRB4*01:01 |
| Ooi__DR15_human | DRB1*15:01,DRB5*01:01 |
| Ooi__DR15_mouse | DRB1*15:01,DRB5*01:01 |
| Ooi__DR1_human | DRB1*01:01 |
| Ooi__DR1_mouse | DRB1*01:01 |
| Racle__3808_HMC | DRB1*03:01,DRB1*07:01,DRB3*01:01,DRB4*01:01,DQA1*02:01-DQB1*02:01,DQA1*02:01-DQB1*02:02,DQA1*05:01-DQB1*02:01,DQA1*05:01-DQB1*02:02,DPA1*01:03-DPB1*03:01,DPA1*01:03-DPB1*11:01,DPA1*02:01-DPB1*03:01,DPA1*02:01-DPB1*11:01 |
| Racle__3808_HMC_DQP | DRB1*03:01,DRB1*07:01,DRB3*01:01,DRB4*01:01,DQA1*02:01-DQB1*02:01,DQA1*02:01-DQB1*02:02,DQA1*05:01-DQB1*02:01,DQA1*05:01-DQB1*02:02,DPA1*01:03-DPB1*03:01,DPA1*01:03-DPB1*11:01,DPA1*02:01-DPB1*03:01,DPA1*02:01-DPB1*11:01 |
| Racle__3808_HMC_DR | DRB1*03:01,DRB1*07:01,DRB3*01:01,DRB4*01:01,DQA1*02:01-DQB1*02:01,DQA1*02:01-DQB1*02:02,DQA1*05:01-DQB1*02:01,DQA1*05:01-DQB1*02:02,DPA1*01:03-DPB1*03:01,DPA1*01:03-DPB1*11:01,DPA1*02:01-DPB1*03:01,DPA1*02:01-DPB1*11:01 |
| Racle__3830NJF | DRB1*04:04,DRB1*11:01,DRB3*02:02,DRB4*01:03,DQA1*03:01-DQB1*03:01,DQA1*03:01-DQB1*03:02,DQA1*05:05-DQB1*03:01,DQA1*05:05-DQB1*03:02,DPA1*01:03-DPB1*02:01,DPA1*01:03-DPB1*06:01 |
| Racle__3830_NJF_DQP | DRB1*04:04,DRB1*11:01,DRB3*02:02,DRB4*01:03,DQA1*03:01-DQB1*03:01,DQA1*03:01-DQB1*03:02,DQA1*05:05-DQB1*03:01,DQA1*05:05-DQB1*03:02,DPA1*01:03-DPB1*02:01,DPA1*01:03-DPB1*06:01 |
| Racle__3830_NJF_DR | DRB1*04:04,DRB1*11:01,DRB3*02:02,DRB4*01:03,DQA1*03:01-DQB1*03:01,DQA1*03:01-DQB1*03:02,DQA1*05:05-DQB1*03:01,DQA1*05:05-DQB1*03:02,DPA1*01:03-DPB1*02:01,DPA1*01:03-DPB1*06:01 |
| Racle__3849BR | DRB1*11:04,DRB3*02:02,DQA1*05:05-DQB1*03:01,DPA1*01:03-DPB1*02:01,DPA1*01:03-DPB1*04:01 |
| Racle__3865DM | DRB1*01:01,DRB1*07:01,DRB4*01:03,DQA1*01:01-DQB1*03:03,DQA1*01:01-DQB1*05:01,DQA1*02:01-DQB1*03:03,DQA1*02:01-DQB1*05:01,DPA1*01:03-DPB1*04:01,DPA1*01:03-DPB1*20:01 |
| Racle__3869_GA | DRB1*01:03,DRB1*04:04,DRB4*01:03,DQA1*03:01-DQB1*03:01,DQA1*03:01-DQB1*03:02,DQA1*05:05-DQB1*03:01,DQA1*05:05-DQB1*03:02,DPA1*01:03-DPB1*04:01,DPA1*01:03-DPB1*12:60 |
| Racle__3869_GA_DQP | DRB1*01:03,DRB1*04:04,DRB4*01:03,DQA1*03:01-DQB1*03:01,DQA1*03:01-DQB1*03:02,DQA1*05:05-DQB1*03:01,DQA1*05:05-DQB1*03:02,DPA1*01:03-DPB1*04:01,DPA1*01:03-DPB1*12:60 |
| Racle__3869_GA_DR | DRB1*01:03,DRB1*04:04,DRB4*01:03,DQA1*03:01-DQB1*03:01,DQA1*03:01-DQB1*03:02,DQA1*05:05-DQB1*03:01,DQA1*05:05-DQB1*03:02,DPA1*01:03-DPB1*04:01,DPA1*01:03-DPB1*12:60 |
| Racle__3911_ME | DRB1*11:01,DRB3*02:02,DQA1*05:05-DQB1*03:01,DPA1*01:03-DPB1*04:01 |
| Racle__3912BAM | DRB1*03:01,DRB1*04:01,DRB3*01:01,DRB4*01:03,DQA1*03:01-DQB1*02:01,DQA1*03:01-DQB1*03:02,DQA1*05:01-DQB1*02:01,DQA1*05:01-DQB1*03:02,DPA1*01:03-DPB1*04:01 |
| Racle__3912_BAM_DQP | DRB1*03:01,DRB1*04:01,DRB3*01:01,DRB4*01:03,DQA1*03:01-DQB1*02:01,DQA1*03:01-DQB1*03:02,DQA1*05:01-DQB1*02:01,DQA1*05:01-DQB1*03:02,DPA1*01:03-DPB1*04:01 |
| Racle__3912_BAM_DR | DRB1*03:01,DRB1*04:01,DRB3*01:01,DRB4*01:03,DQA1*03:01-DQB1*02:01,DQA1*03:01-DQB1*03:02,DQA1*05:01-DQB1*02:01,DQA1*05:01-DQB1*03:02,DPA1*01:03-DPB1*04:01 |
| Racle__3947_GA | DRB1*01:01,DRB1*13:01,DRB3*01:01,DQA1*01:01-DQB1*05:01,DQA1*01:01-DQB1*06:03,DQA1*01:03-DQB1*05:01,DQA1*01:03-DQB1*06:03,DPA1*01:03-DPB1*02:01,DPA1*01:03-DPB1*04:02 |
| Racle__3947_GA_DQP | DRB1*01:01,DRB1*13:01,DRB3*01:01,DQA1*01:01-DQB1*05:01,DQA1*01:01-DQB1*06:03,DQA1*01:03-DQB1*05:01,DQA1*01:03-DQB1*06:03,DPA1*01:03-DPB1*02:01,DPA1*01:03-DPB1*04:02 |
| Racle__3947_GA_DR | DRB1*01:01,DRB1*13:01,DRB3*01:01,DQA1*01:01-DQB1*05:01,DQA1*01:01-DQB1*06:03,DQA1*01:03-DQB1*05:01,DQA1*01:03-DQB1*06:03,DPA1*01:03-DPB1*02:01,DPA1*01:03-DPB1*04:02 |
| Racle__3971_ORA | DRB1*13:03,DRB1*07:01,DRB3*01:01,DRB4*01:01,DQA1*02:01-DQB1*02:02,DQA1*02:01-DQB1*03:01,DQA1*05:05-DQB1*02:02,DQA1*05:05-DQB1*03:01,DPA1*01:03-DPB1*04:01,DPA1*02:02-DPB1*04:01 |
| Racle__3971_ORA_DR | DRB1*13:03,DRB1*07:01,DRB3*01:01,DRB4*01:01,DQA1*02:01-DQB1*02:02,DQA1*02:01- |
|  | DQB1*03:01,DQA1*05:05-DQB1*02:02,DQA1*05:05-DQB1*03:01,DPA1*01:03-DPB1*04:01,DPA1*02:02-DPB1*04:01 |
| Racle__3993 | DRB1*07:01,DRB1*15:01,DRB4*01:03,DRB5*01:01,DQA1*01:02-DQB1*02:02,DQA1*01:02-DQB1*06:02,DQA1*02:01-DQB1*02:02,DQA1*02:01-DQB1*06:02,DPA1*01:03-DPB1*04:01,DPA1*01:03-DPB1*17:01,DPA1*02:01-DPB1*04:01,DPA1*02:01-DPB1*17:01 |
| Racle__4001 | DRB1*13:01,DRB1*14:01,DRB3*01:01,DRB3*02:02,DQA1*01:03-DQB1*05:03,DQA1*01:03-DQB1*06:03,DQA1*01:04-DQB1*05:03,DQA1*01:04-DQB1*06:03,DPA1*01:03-DPB1*04:01,DPA1*01:03-DPB1*04:02 |
| Racle__4001_DR | DRB1*13:01,DRB1*14:01,DRB3*01:01,DRB3*02:02,DQA1*01:03-DQB1*05:03,DQA1*01:03-DQB1*06:03,DQA1*01:04-DQB1*05:03,DQA1*01:04-DQB1*06:03,DPA1*01:03-DPB1*04:01,DPA1*01:03-DPB1*04:02 |
| Racle__4021 | DRB1*11:01,DRB1*04:05,DRB3*02:02,DRB4*01:03,DQA1*03:03-DQB1*02:02,DQA1*03:03-DQB1*03:01,DQA1*05:05-DQB1*02:02,DQA1*05:05-DQB1*03:01,DPA1*01:03-DPB1*03:01,DPA1*01:03-DPB1*10:40 |
| Racle__4021_DQP | DRB1*11:01,DRB1*04:05,DRB3*02:02,DRB4*01:03,DQA1*03:03-DQB1*02:02,DQA1*03:03-DQB1*03:01,DQA1*05:05-DQB1*02:02,DQA1*05:05-DQB1*03:01,DPA1*01:03-DPB1*03:01,DPA1*01:03-DPB1*10:40 |
| Racle__4021_DR | DRB1*11:01,DRB1*04:05,DRB3*02:02,DRB4*01:03,DQA1*03:03-DQB1*02:02,DQA1*03:03-DQB1*03:01,DQA1*05:05-DQB1*02:02,DQA1*05:05-DQB1*03:01,DPA1*01:03-DPB1*03:01,DPA1*01:03-DPB1*10:40 |
| Racle__4037_DC | DRB1*01:01,DQA1*01:01-DQB1*05:01,DPA1*01:03-DPB1*04:01,DPA1*01:03-DPB1*06:01 |
| Racle__4052_BA | DRB1*03:01,DRB1*11:04,DRB3*01:01,DRB3*02:02,DQA1*05:01-DQB1*02:01,DQA1*05:01-DQB1*03:01,DQA1*05:05-DQB1*02:01,DQA1*05:05-DQB1*03:01,DPA1*01:03-DPB1*04:01 |
| Racle__4052_BA_DQP | DRB1*03:01,DRB1*11:04,DRB3*01:01,DRB3*02:02,DQA1*05:01-DQB1*02:01,DQA1*05:01-DQB1*03:01,DQA1*05:05-DQB1*02:01,DQA1*05:05-DQB1*03:01,DPA1*01:03-DPB1*04:01 |
| Racle__4052_BA_DR | DRB1*03:01,DRB1*11:04,DRB3*01:01,DRB3*02:02,DQA1*05:01-DQB1*02:01,DQA1*05:01-DQB1*03:01,DQA1*05:05-DQB1*02:01,DQA1*05:05-DQB1*03:01,DPA1*01:03-DPB1*04:01 |
| Racle__BP455 | DRB1*10:01,DRB1*13:01,DRB3*01:01,DQA1*01:05-DQB1*05:01,DQA1*01:05-DQB1*06:03,DQA1*01:10-DQB1*05:01,DQA1*01:10-DQB1*06:03,DPA1*01:03-DPB1*02:01 |
| Racle__CD165 | DRB1*11:01,DRB3*02:02,DQA1*05:05-DQB1*03:01,DPA1*01:03-DPB1*04:01,DPA1*01:03-DPB1*04:02 |
| Racle__CD165_DQP | DRB1*11:01,DRB3*02:02,DQA1*05:05-DQB1*03:01,DPA1*01:03-DPB1*04:01,DPA1*01:03-DPB1*04:02 |
| Racle__CD165_DR | DRB1*11:01,DRB3*02:02,DQA1*05:05-DQB1*03:01,DPA1*01:03-DPB1*04:01,DPA1*01:03-DPB1*04:02 |
| Racle__CM647 | DRB1*07:01,DRB1*16:01,DRB4*01:03,DRB5*02:02,DQA1*01:02-DQB1*02:02,DQA1*01:02-DQB1*05:02,DQA1*02:01-DQB1*02:02,DQA1*02:01-DQB1*05:02,DPA1*01:03-DPB1*02:01,DPA1*01:03-DPB1*23:01 |
| Racle__CM647_DQP | DRB1*07:01,DRB1*16:01,DRB4*01:03,DRB5*02:02,DQA1*01:02-DQB1*02:02,DQA1*01:02-DQB1*05:02,DQA1*02:01-DQB1*02:02,DQA1*02:01-DQB1*05:02,DPA1*01:03-DPB1*02:01,DPA1*01:03-DPB1*23:01 |
| Racle__CM647_DR | DRB1*07:01,DRB1*16:01,DRB4*01:03,DRB5*02:02,DQA1*01:02-DQB1*02:02,DQA1*01:02-DQB1*05:02,DQA1*02:01-DQB1*02:02,DQA1*02:01-DQB1*05:02,DPA1*01:03-DPB1*02:01,DPA1*01:03-DPB1*23:01 |
| Racle__GD149 | DRB1*07:01,DRB1*13:01,DRB3*01:01,DRB4*01:01,DQA1*01:10-DQB1*02:02,DQA1*01:10-DQB1*06:03,DQA1*02:01-DQB1*02:02,DQA1*02:01-DQB1*06:03,DPA1*01:03-DPB1*03:01,DPA1*01:03-DPB1*04:01,DPA1*02:01-DPB1*03:01,DPA1*02:01-DPB1*04:01 |
| Racle__JY | DRB1*04:04,DRB1*13:01,DRB3*01:01,DRB4*01:03,DQA1*01:03-DQB1*03:02,DQA1*01:03-DQB1*06:03,DQA1*03:01-DQB1*03:02,DQA1*03:01-DQB1*06:03,DPA1*01:03-DPB1*02:01,DPA1*01:03-DPB1*04:01 |
| Racle__JY_DR | DRB1*04:04,DRB1*13:01,DRB3*01:01,DRB4*01:03,DQA1*01:03-DQB1*03:02,DQA1*01:03-DQB1*06:03,DQA1*03:01-DQB1*03:02,DQA1*03:01-DQB1*06:03,DPA1*01:03-DPB1*02:01,DPA1*01:03-DPB1*04:01 |
| Racle__PD42 | DRB1*01:02,DRB1*15:01,DRB5*01:01,DQA1*01:01-DQB1*05:01,DQA1*01:01-DQB1*06:02,DQA1*01:02-DQB1*05:01,DQA1*01:02-DQB1*06:02,DPA1*01:03-DPB1*04:01,DPA1*01:03-DPB1*05:01,DPA1*02:02-DPB1*04:01,DPA1*02:02-DPB1*05:01 |
| Racle__RA957 | DRB1*04:01,DRB1*08:01,DRB4*01:03,DQA1*03:03-DQB1*03:01,DQA1*03:03-DQB1*04:02,DQA1*04:01-DQB1*03:01,DQA1*04:01-DQB1*04:02,DPA1*01:03-DPB1*04:01,DPA1*01:03-DPB1*04:02 |
| Racle__RA957_DQP | DRB1*04:01,DRB1*08:01,DRB4*01:03,DQA1*03:03-DQB1*03:01,DQA1*03:03-DQB1*04:02,DQA1*04:01-DQB1*03:01,DQA1*04:01-DQB1*04:02,DPA1*01:03-DPB1*04:01,DPA1*01:03-DPB1*04:02 |
| Racle__RA957_DR | DRB1*04:01,DRB1*08:01,DRB4*01:03,DQA1*03:03-DQB1*03:01,DQA1*03:03-DQB1*04:02,DQA1*04:01-DQB1*03:01,DQA1*04:01-DQB1*04:02,DPA1*01:03-DPB1*04:01,DPA1*01:03-DPB1*04:02 |
| Racle__TIL1 | DRB1*01:01,DRB1*04:08,DRB4*01:03,DQA1*01:01-DQB1*03:01,DQA1*01:01-DQB1*05:01,DQA1*03:03-DQB1*03:01,DQA1*03:03-DQB1*05:01,DPA1*01:03-DPB1*02:01,DPA1*01:03-DPB1*04:01 |
| Racle__TIL3 | DRB1*12:01,DRB1*15:01,DRB3*02:02,DRB5*01:01,DQA1*01:02-DQB1*03:01,DQA1*01:02-DQB1*05:02,DQA1*05:05-DQB1*03:01,DQA1*05:05-DQB1*05:02,DPA1*01:03-DPB1*03:01,DPA1*01:03-DPB1*04:01 |
| Ritz__DOHH2_DQ | DRB1*01:01,DRB1*15:01,DRB5*01:01,DQA1*01:01-DQB1*05:01,DQA1*01:02-DQB1*06:02,DQA1*01:02-DQB1*05:01,DQA1*01:01-DQB1*06:02 |
| Ritz__DOHH2_DR | DRB1*01:01,DRB1*15:01,DRB5*01:01,DQA1*01:01-DQB1*05:01,DQA1*01:02-DQB1*06:02,DQA1*01:02-DQB1*05:01,DQA1*01:01-DQB1*06:02 |
| Ritz__Maver_1_DQ | DRB1*01:01,DRB1*13:01,DRB3*02:02,DQA1*01:01-DQB1*05:01,DQA1*01:03-DQB1*06:03,DQA1*01:01-DQB1*06:03,DQA1*01:03-DQB1*05:01 |
| Ritz__Maver_1_DR | DRB1*01:01,DRB1*13:01,DRB3*02:02,DQA1*01:01-DQB1*05:01,DQA1*01:03-DQB1*06:03,DQA1*01:01-DQB1*06:03,DQA1*01:03-DQB1*05:01 |
| Saghar__peptide_0901 | DRB1*09:01,DRB4*01:03 |
| Saghar__peptide_1401 | DRB1*14:01,DRB3*02:02 |
| Saghar__peptide_1601 | DRB1*16:01,DRB5*02:02 |
| Saghar__peptide_HeLa | DRB1*01:02 |
| Scally__DRB1_0401 | DRB1*04:01,DRB4*01:01 |
| Scally__DRB1_0402 | DRB1*04:02 |
| Scally__DRB1_0404 | DRB1*04:04,DRB4*01:01 |
| Sofron__A20 | H-2-IAd,H-2-IEd |
| Sofron__Balb_c_1 | H-2-IAd,H-2-IEd |
| Sofron__Balb_c_2 | H-2-IAd,H-2-IEd |
| Sofron__C57BL_6_1 | H-2-IAb |
| Sofron__C57BL_6_2 | H-2-IAb |
| Ting__T2_DRB1_0405 | DRB1*04:05,DRB4*01:01 |
| Wang__LA3 | DRB1*03:01,DRB1*15:01,DRB3*01:01,DRB5*01:01 |
| Wang__LA4 | DRB1*11:01,DRB1*13:02,DRB3*02:02,DRB3*03:01 |
| Wang__LA5 | DRB1*03:01,DRB1*03:05,DRB3*01:01 |
| Wang__LA7 | DRB1*04:03,DRB1*15:01,DRB5*01:01 |
| Wang__LA8 | DRB1*04:01,DRB1*10:01,DRB4*01:01 |
| Wang__RA2 | DRB1*01:01,DRB1*04:01,DRB4*01:01 |
| Wang__RA3 | DRB1*01:01,DRB1*04:01,DRB4*01:01 |

**Supplementary table 2:** Novel peptide ligand dataset, in the format PEPTIDE 1 CELL_LINE_ID CONTEXT. The data can be found in the external datafile *saghar_peptide_data.xlsx*.

**Supplementary table 3:** Migration of peptides annotated towards HLA-DQ by the models trained with (w_Saghar) and without (wo_Saghar) the novel data.

| Peptides uniquely assigned to HLA-DQ in w_Saghar method | Distribution of peptide annotations in wo_Saghar method |  |  |
| --- | --- | --- | --- |
| 9309 | HLA-DP<br>1816 | HLA-DR<br>4688 | Trash<br>2805 |
| Peptides uniquely assigned to HLA-DQ in wo_Saghar method | Distribution of peptide annotations in w_Saghar method |  |  |
| 4316 | HLA-DP<br>609 | HLA-DR<br>2629 | Trash<br>1078 |

**Supplementary table 4:** Overview of peptides assigned to DQ molecules in the methods with (w_Saghar) and without (wo_Saghar) the novel data. Trash peptides with percentile rank greater than 20 are not included in the metrics. A bold value indicates either a higher average peptide count or a lower mean/median percentile rank in a given method.

| Molecule | Average number of peptides per dataset |  | Average of per-dataset mean percentile rank |  | Median of per-dataset median percentile rank |  |
| --- | --- | --- | --- | --- | --- | --- |
|  | w_Saghar | wo_Saghar | w_Saghar | wo_Saghar | w_Saghar | wo_Saghar |
| HLA-DQA1*01:01-DQB1*02:01 | 6.00 | **7.50** | **3.2818** | 3.7402 | **0.6713** | 0.8226 |
| HLA-DQA1*01:01-DQB1*03:01 | 20.00 | **170.00** | **0.3084** | 1.8631 | **0.0086** | 0.4087 |
| HLA-DQA1*01:01-DQB1*03:03 | 27.00 | **58.00** | **0.4281** | 0.5240 | **0.0078** | 0.0086 |
| HLA-DQA1*01:01-DQB1*05:01 | **209.53** | 155.06 | 2.6547 | **1.1562** | 1.1783 | **0.1096** |
| HLA-DQA1*01:01-DQB1*06:02 | **30.50** | 27.67 | 1.1216 | **0.9356** | **0.2011** | 0.2064 |
| HLA-DQA1*01:01-DQB1*06:03 | 26.40 | **59.20** | **1.2573** | 2.6273 | **0.1503** | 2.1822 |
| HLA-DQA1*01:02-DQB1*02:02 | 13.00 | **28.75** | **0.1723** | 1.3248 | **0.0073** | 0.0094 |
| HLA-DQA1*01:02-DQB1*03:01 | 53.00 | **99.00** | **0.3111** | 1.5251 | **0.0083** | 0.0552 |
| HLA-DQA1*01:02-DQB1*03:02 | 6.50 | **8.50** | **0.3849** | 0.8472 | **0.0073** | 0.0078 |
| HLA-DQA1*01:02-DQB1*05:01 | 27.33 | **43.33** | 2.2276 | **0.8075** | 0.1011 | **0.0092** |
| HLA-DQA1*01:02-DQB1*05:02 | **512.20** | 368.00 | 2.7410 | **1.1782** | 0.6769 | **0.0513** |
| HLA-DQA1*01:02-DQB1*06:02 | 550.88 | **551.88** | 4.1209 | **3.8584** | 1.9511 | **1.5487** |
| HLA-DQA1*01:02-DQB1*06:04 | **1548.50** | 1520.00 | **1.8068** | 1.9438 | **0.3963** | 0.4895 |
| HLA-DQA1*01:03-DQB1*03:02 | 23.50 | **54.50** | **0.3737** | 0.9327 | **0.0071** | 0.0091 |
| HLA-DQA1*01:03-DQB1*05:01 | 6.60 | **12.80** | 0.6552 | **0.3573** | **0.0140** | 0.0165 |
| HLA-DQA1*01:03-DQB1*05:03 | **2.00** | 1.50 | 4.2220 | **0.0102** | 4.2220 | **0.0102** |
| HLA-DQA1*01:03-DQB1*06:03 | **353.10** | 325.60 | **2.6588** | 3.0594 | **0.5745** | 0.5782 |
| HLA-DQA1*01:04-DQB1*05:03 | **549.00** | 406.33 | 2.8193 | **0.7154** | **1.7952** | 0.2739 |
| HLA-DQA1*01:04-DQB1*06:03 | 3.00 | **3.50** | **3.2792** | 3.5277 | 4.1573 | **2.3288** |
| HLA-DQA1*01:05-DQB1*05:01 | **89.00** | 27.00 | 2.5292 | **1.4787** | 1.2150 | **0.0631** |
| HLA-DQA1*01:05-DQB1*06:03 | 27.00 | **51.00** | **0.6273** | 0.9981 | **0.0100** | 0.0225 |
| HLA-DQA1*01:10-DQB1*02:02 | 6.00 | **32.00** | **0.0063** | 0.0663 | **0.0067** | 0.0073 |
| HLA-DQA1*01:10-DQB1*05:01 | 17.00 | **22.00** | **0.0983** | 0.9877 | **0.0077** | 0.0085 |
| HLA-DQA1*01:10-DQB1*06:03 | **205.00** | 189.00 | **1.9616** | 1.9742 | 0.5409 | **0.4139** |
| HLA-DQA1*02:01-DQB1*02:01 | **75.67** | 73.33 | **2.2314** | 2.3259 | 0.5515 | **0.3520** |
| HLA-DQA1*02:01-DQB1*02:02 | 1188.91 | **1207.82** | 3.4357 | **2.9958** | 1.1275 | **1.1813** |
| HLA-DQA1*02:01-DQB1*03:01 | 8.00 | **10.50** | 6.1621 | **4.9487** | **2.2676** | 2.6416 |
| HLA-DQA1*02:01-DQB1*03:03 | **868.00** | 711.00 | 3.8598 | **2.8138** | 2.0423 | **0.9332** |
| HLA-DQA1*02:01-DQB1*05:01 | 11.00 | **24.00** | **0.0094** | 1.5804 | **0.0060** | 0.0080 |
| HLA-DQA1*02:01-DQB1*05:02 | **18.33** | 15.67 | **0.0088** | 2.0617 | **0.0059** | 0.0083 |
| HLA-DQA1*02:01-DQB1*06:02 | 6.00 | **7.00** | 1.0585 | **0.0980** | 0.0355 | **0.0091** |
| HLA-DQA1*02:01-DQB1*06:03 | 15.00 | 15.00 | **0.0088** | 0.2958 | 0.0085 | **0.0072** |
| HLA-DQA1*03:01-DQB1*02:01 | **14.67** | 14.33 | 3.0440 | **2.0955** | **0.0953** | 0.9899 |
| HLA-DQA1*03:01-DQB1*03:01 | **54.67** | 16.83 | 3.5246 | **0.7732** | 1.1628 | **0.0092** |
| HLA-DQA1*03:01-DQB1*03:02 | **152.86** | 92.64 | 4.2735 | **1.5354** | 2.0231 | **0.1016** |
| HLA-DQA1*03:01-DQB1*06:02 | **4.00** | 1.00 | 2.3313 | **0.4498** | **0.2132** | 0.4498 |
| HLA-DQA1*03:01-DQB1*06:03 | 8.00 | **10.00** | 0.6198 | **0.0115** | 0.0270 | **0.0088** |
| HLA-DQA1*03:02-DQB1*03:03 | **1565.00** | 1003.00 | 3.3282 | **2.6386** | 1.4368 | **0.6301** |
| HLA-DQA1*03:03-DQB1*02:02 | **48.33** | 15.00 | 0.8371 | **0.1817** | **0.0096** | 0.0109 |
| HLA-DQA1*03:03-DQB1*03:01 | **593.25** | 389.50 | 3.0677 | **1.0865** | 0.8015 | **0.0512** |
| HLA-DQA1*03:03-DQB1*04:02 | **82.00** | 17.00 | **0.8113** | 1.2414 | **0.0109** | 0.0204 |
| HLA-DQA1*03:03-DQB1*05:01 | **9.00** | 4.00 | **0.0143** | 0.0253 | **0.0085** | 0.0258 |
| HLA-DQA1*04:01-DQB1*03:01 | 96.67 | **371.00** | **1.4585** | 3.5739 | **0.0179** | 1.5131 |
| HLA-DQA1*04:01-DQB1*04:02 | **967.60** | 801.80 | 3.0511 | **2.3695** | 1.4733 | **0.5541** |
| HLA-DQA1*05:01-DQB1*02:01 | 344.00 | **347.79** | **2.2603** | 2.6852 | **0.2518** | 0.3707 |
| HLA-DQA1*05:01-DQB1*03:01 | 1008.50 | 1008.50 | 4.6531 | **4.3624** | 1.7700 | **1.6834** |
| HLA-DQA1*05:01-DQB1*03:02 | 19.67 | **22.00** | 3.5080 | **2.4423** | **0.2787** | 0.5770 |
| HLA-DQA1*05:01-DQB1*05:01 | 2.00 | **4.00** | **0.0071** | 0.0238 | **0.0070** | 0.0082 |
| HLA-DQA1*05:05-DQB1*02:02 | **64.00** | 62.40 | 2.4021 | **2.1639** | **0.1911** | 0.3171 |
| HLA-DQA1*05:05-DQB1*03:01 | 577.55 | **594.90** | **3.3196** | 3.8926 | **1.4065** | 1.8265 |
| HLA-DQA1*05:05-DQB1*03:02 | 18.67 | **25.33** | **1.2711** | 1.5989 | 0.0884 | **0.0495** |
| HLA-DQA1*05:05-DQB1*05:02 | **24.00** | 9.00 | 0.0093 | **0.0068** | **0.0064** | 0.0071 |

**Supplementary table 5:** Overview of consistency analysis in the methods with (w_Saghar) and without (wo_Saghar) the novel data. The molecules are sorted in descending order by the difference in mean consistency. Further, the metrics are calculated on the peptide sets used in the consistency analysis, with the union of identified trash peptides removed. As such, the metrics regarding differences in percentile ranks may not correspond one-to-one with the values in supplementary table 4.

| Molecules | Difference in mean consistency (w_Saghar - wo_Saghar) | Number of cell lines with at least 20 peptides in both methods | Number of cell lines with larger peptide count in w_Saghar | Number of cell lines with lower mean %rank in w_Saghar | Fraction of cell lines with either larger peptide count or lower mean %rank in w_Saghar |
| --- | --- | --- | --- | --- | --- |
| HLA-DQA1*03:01-DQB1*03:02 | 0.148 | 6 | 6 | 1 | 1.000 |
| HLA-DQA1*03:01-DQB1*03:01 | 0.147 | 2 | 2 | 0 | 1.000 |
| HLA-DQA1*03:03-DQB1*03:01 | 0.119 | 5 | 5 | 1 | 1.000 |
| HLA-DQA1*01:01-DQB1*05:01 | 0.079 | 10 | 10 | 3 | 1.000 |
| HLA-DQA1*04:01-DQB1*04:02 | 0.039 | 5 | 3 | 2 | 0.600 |
| HLA-DQA1*01:02-DQB1*05:02 | 0.037 | 5 | 5 | 1 | 1.000 |
| HLA-DQA1*02:01-DQB1*03:03 | 0.035 | 2 | 2 | 1 | 1.000 |
| HLA-DQA1*01:10-DQB1*06:03 | 0.034 | 2 | 2 | 1 | 1.000 |
| HLA-DQA1*05:05-DQB1*03:01 | 0.027 | 16 | 2 | 8 | 0.625 |
| HLA-DQA1*01:02-DQB1*06:02 | 0.020 | 7 | 3 | 1 | 0.429 |
| HLA-DQA1*02:01-DQB1*02:01 | 0.017 | 3 | 2 | 1 | 0.667 |
| HLA-DQA1*05:01-DQB1*02:01 | 0.016 | 12 | 4 | 9 | 0.917 |
| HLA-DQA1*05:05-DQB1*02:02 | 0.014 | 3 | 0 | 2 | 0.667 |
| HLA-DQA1*02:01-DQB1*02:02 | 0.010 | 9 | 1 | 8 | 1.000 |
| HLA-DQA1*05:05-DQB1*03:02 | 0.005 | 2 | 1 | 1 | 1.000 |
| HLA-DQA1*05:01-DQB1*03:01 | 0.002 | 4 | 3 | 0 | 0.750 |
| HLA-DQA1*01:02-DQB1*06:04 | -0.002 | 2 | 1 | 2 | 1.000 |
| HLA-DQA1*01:03-DQB1*03:02 | -0.008 | 2 | 0 | 2 | 1.000 |
| HLA-DQA1*01:03-DQB1*06:03 | -0.009 | 7 | 5 | 4 | 0.857 |
| HLA-DQA1*01:01-DQB1*06:03 | -0.027 | 2 | 0 | 2 | 1.000 |
| HLA-DQA1*04:01-DQB1*03:01 | -0.031 | 3 | 0 | 3 | 1.000 |

**Supplementary table 6:** Overview of DQ-heterozygous datasets used in the cis vs trans-only DQ analysis, along with their DQ HLA typing. Molecules marked in red are trans-only heterodimers. Molecules in bold are part of the DQ-SA training data.

| Dataset | DQ heterodimers |  |  |  |
| --- | --- | --- | --- | --- |
| Racle__3808_HMC | DQA1*02:01-DQB1*02:01 | **DQA1\*02:01-DQB1\*02:02** | **DQA1\*05:01-DQB1\*02:01** | DQA1*05:01-DQB1*02:02 |
| Racle__3830NJF | DQA1*03:01-DQB1*03:01 | **DQA1\*03:01-DQB1\*03:02** | **DQA1\*05:05-DQB1\*03:01** | DQA1*05:05-DQB1*03:02 |
| Racle__3830_NJF_DQP | DQA1*03:01-DQB1*03:01 | **DQA1\*03:01-DQB1\*03:02** | **DQA1\*05:05-DQB1\*03:01** | DQA1*05:05-DQB1*03:02 |
| Racle__3865DM | **DQA1\*01:01-DQB1\*03:03** | **DQA1\*01:01-DQB1\*05:01** | **DQA1\*02:01-DQB1\*03:03** | **DQA1\*02:01-DQB1\*05:01** |
| Racle__3869_GA | DQA1*03:01-DQB1*03:01 | **DQA1\*03:01-DQB1\*03:02** | **DQA1\*05:05-DQB1\*03:01** | DQA1*05:05-DQB1*03:02 |
| Racle__3869_GA_DQP | DQA1*03:01-DQB1*03:01 | **DQA1\*03:01-DQB1\*03:02** | **DQA1\*05:05-DQB1\*03:01** | DQA1*05:05-DQB1*03:02 |
| Racle__3912BAM | DQA1*03:01-DQB1*02:01 | **DQA1\*03:01-DQB1\*03:02** | **DQA1\*05:01-DQB1\*02:01** | DQA1*05:01-DQB1*03:02 |
| Racle__3912_BAM_DR | DQA1*03:01-DQB1*02:01 | **DQA1\*03:01-DQB1\*03:02** | **DQA1\*05:01-DQB1\*02:01** | DQA1*05:01-DQB1*03:02 |
| Racle__3947_GA | **DQA1\*01:01-DQB1\*05:01** | DQA1*01:01-DQB1*06:03 | DQA1*01:03-DQB1*05:01 | **DQA1\*01:03-DQB1\*06:03** |
| Racle__4021 | DQA1*03:03-DQB1*02:02 | **DQA1\*03:03-DQB1\*03:01** | DQA1*05:05-DQB1*02:02 | **DQA1\*05:05-DQB1\*03:01** |
| Racle__4021_DQP | DQA1*03:03-DQB1*02:02 | **DQA1\*03:03-DQB1\*03:01** | DQA1*05:05-DQB1*02:02 | **DQA1\*05:05-DQB1\*03:01** |
| Racle__4052_BA | **DQA1\*05:01-DQB1\*02:01** | DQA1*05:01-DQB1*03:01 | DQA1*05:05-DQB1*02:01 | **DQA1\*05:05-DQB1\*03:01** |
| Racle__4052_BA_DQP | **DQA1\*05:01-DQB1\*02:01** | DQA1*05:01-DQB1*03:01 | DQA1*05:05-DQB1*02:01 | **DQA1\*05:05-DQB1\*03:01** |
| Racle__BP455 | DQA1*01:05-DQB1*05:01 | DQA1*01:05-DQB1*06:03 | DQA1*01:10-DQB1*05:01 | DQA1*01:10-DQB1*06:03 |
| Racle__CM647 | **DQA1\*01:02-DQB1\*02:02** | **DQA1\*01:02-DQB1\*05:02** | **DQA1\*02:01-DQB1\*02:02** | **DQA1\*02:01-DQB1\*05:02** |
| Racle__CM647_DQP | **DQA1\*01:02-DQB1\*02:02** | **DQA1\*01:02-DQB1\*05:02** | **DQA1\*02:01-DQB1\*02:02** | **DQA1\*02:01-DQB1\*05:02** |
| Racle__CM647_DR | **DQA1\*01:02-DQB1\*02:02** | **DQA1\*01:02-DQB1\*05:02** | **DQA1\*02:01-DQB1\*02:02** | **DQA1\*02:01-DQB1\*05:02** |
| Racle__GD149 | **DQA1\*01:10-DQB1\*02:02** | DQA1*01:10-DQB1*06:03 | **DQA1\*02:01-DQB1\*02:02** | **DQA1\*02:01-DQB1\*06:03** |
| Racle__JY | **DQA1\*01:03-DQB1\*03:02** | **DQA1\*01:03-DQB1\*06:03** | **DQA1\*03:01-DQB1\*03:02** | **DQA1\*03:01-DQB1\*06:03** |
| Racle__JY_DR | **DQA1\*01:03-DQB1\*03:02** | **DQA1\*01:03-DQB1\*06:03** | **DQA1\*03:01-DQB1\*03:02** | **DQA1\*03:01-DQB1\*06:03** |
| Racle__PD42 | **DQA1\*01:01-DQB1\*05:01** | DQA1*01:01-DQB1*06:02 | DQA1*01:02-DQB1*05:01 | **DQA1\*01:02-DQB1\*06:02** |
| Racle__RA957 | **DQA1\*03:03-DQB1\*03:01** | DQA1*03:03-DQB1*04:02 | DQA1*04:01-DQB1*03:01 | **DQA1\*04:01-DQB1\*04:02** |
| Racle__RA957_DQP | **DQA1\*03:03-DQB1\*03:01** | DQA1*03:03-DQB1*04:02 | DQA1*04:01-DQB1*03:01 | **DQA1\*04:01-DQB1\*04:02** |
| Racle__RA957_DR | **DQA1\*03:03-DQB1\*03:01** | DQA1*03:03-DQB1*04:02 | DQA1*04:01-DQB1*03:01 | **DQA1\*04:01-DQB1\*04:02** |
| Racle__TIL1 | **DQA1\*01:01-DQB1\*03:01** | **DQA1\*01:01-DQB1\*05:01** | **DQA1\*03:03-DQB1\*03:01** | **DQA1\*03:03-DQB1\*05:01** |
| Racle__TIL3 | **DQA1\*01:02-DQB1\*03:01** | **DQA1\*01:02-DQB1\*05:02** | **DQA1\*05:05-DQB1\*03:01** | **DQA1\*05:05-DQB1\*05:02** |
| Ritz__DOHH2_DQ | **DQA1\*01:01-DQB1\*05:01** | **DQA1\*01:02-DQB1\*06:02** | DQA1*01:02-DQB1*05:01 | DQA1*01:01-DQB1*06:02 |
| Ritz__Maver_1_DQ | **DQA1\*01:01-DQB1\*05:01** | **DQA1\*01:03-DQB1\*06:03** | DQA1*01:01-DQB1*06:03 | DQA1*01:03-DQB1*05:01 |

**Supplementary table 7:** DQ molecules covered by the method including the novel data. The highlighted molecules are covered by a peptide count of at least 100, while the remaining molecules have a distance of at most 0.025 to one of the highlighted molecules.

|  |  |  |  |
| --- | --- | --- | --- |
| HLA-DQA1*01:01-DQB1*05:01 | HLA-DQA1*01:01-DQB1*05:02 | HLA-DQA1*03:01-DQB1*02:02 | HLA-DQA1*05:01-DQB1*03:19 |
| HLA-DQA1*01:01-DQB1*06:03 | HLA-DQA1*01:01-DQB1*05:03 | HLA-DQA1*03:01-DQB1*03:03 | HLA-DQA1*05:01-DQB1*03:22 |
| HLA-DQA1*01:02-DQB1*05:02 | HLA-DQA1*01:01-DQB1*06:02 | HLA-DQA1*03:01-DQB1*03:19 | HLA-DQA1*05:01-DQB1*03:62 |
| HLA-DQA1*01:02-DQB1*06:02 | HLA-DQA1*01:01-DQB1*06:04 | HLA-DQA1*03:01-DQB1*03:22 | HLA-DQA1*05:02-DQB1*02:01 |
| HLA-DQA1*01:02-DQB1*06:04 | HLA-DQA1*01:02-DQB1*05:01 | HLA-DQA1*03:01-DQB1*03:62 | HLA-DQA1*05:02-DQB1*02:02 |
| HLA-DQA1*01:03-DQB1*06:03 | HLA-DQA1*01:02-DQB1*05:03 | HLA-DQA1*03:01-DQB1*03:87 | HLA-DQA1*05:02-DQB1*03:01 |
| HLA-DQA1*01:04-DQB1*05:03 | HLA-DQA1*01:02-DQB1*05:46 | HLA-DQA1*03:01-DQB1*04:01 | HLA-DQA1*05:02-DQB1*03:02 |
| HLA-DQA1*01:10-DQB1*06:03 | HLA-DQA1*01:02-DQB1*06:03 | HLA-DQA1*03:01-DQB1*04:02 | HLA-DQA1*05:03-DQB1*02:01 |
| HLA-DQA1*02:01-DQB1*02:01 | HLA-DQA1*01:03-DQB1*05:02 | HLA-DQA1*03:02-DQB1*02:01 | HLA-DQA1*05:03-DQB1*02:02 |
| HLA-DQA1*02:01-DQB1*02:02 | HLA-DQA1*01:03-DQB1*06:02 | HLA-DQA1*03:02-DQB1*02:02 | HLA-DQA1*05:03-DQB1*03:01 |
| HLA-DQA1*02:01-DQB1*03:03 | HLA-DQA1*01:03-DQB1*06:04 | HLA-DQA1*03:02-DQB1*03:01 | HLA-DQA1*05:03-DQB1*03:02 |
| HLA-DQA1*03:01-DQB1*03:01 | HLA-DQA1*01:04-DQB1*05:01 | HLA-DQA1*03:02-DQB1*03:02 | HLA-DQA1*05:03-DQB1*03:22 |
| HLA-DQA1*03:01-DQB1*03:02 | HLA-DQA1*01:04-DQB1*05:02 | HLA-DQA1*03:02-DQB1*03:22 | HLA-DQA1*05:05-DQB1*02:01 |
| HLA-DQA1*03:02-DQB1*03:03 | HLA-DQA1*01:04-DQB1*06:02 | HLA-DQA1*03:02-DQB1*04:01 | HLA-DQA1*05:05-DQB1*03:22 |
| HLA-DQA1*03:03-DQB1*02:02 | HLA-DQA1*01:04-DQB1*06:03 | HLA-DQA1*03:02-DQB1*04:02 | HLA-DQA1*05:06-DQB1*03:01 |
| HLA-DQA1*03:03-DQB1*03:01 | HLA-DQA1*01:04-DQB1*06:04 | HLA-DQA1*03:03-DQB1*02:01 | HLA-DQA1*05:06-DQB1*03:02 |
| HLA-DQA1*03:03-DQB1*04:02 | HLA-DQA1*01:05-DQB1*05:01 | HLA-DQA1*03:03-DQB1*03:02 | HLA-DQA1*05:08-DQB1*02:01 |
| HLA-DQA1*04:01-DQB1*03:01 | HLA-DQA1*01:05-DQB1*05:02 | HLA-DQA1*03:03-DQB1*03:03 | HLA-DQA1*05:08-DQB1*03:01 |
| HLA-DQA1*04:01-DQB1*04:02 | HLA-DQA1*01:05-DQB1*05:03 | HLA-DQA1*03:03-DQB1*03:22 | HLA-DQA1*05:08-DQB1*03:02 |
| HLA-DQA1*05:01-DQB1*02:01 | HLA-DQA1*01:05-DQB1*06:02 | HLA-DQA1*03:03-DQB1*04:01 | HLA-DQA1*06:01-DQB1*03:01 |
| HLA-DQA1*05:01-DQB1*03:01 | HLA-DQA1*01:05-DQB1*06:03 | HLA-DQA1*04:01-DQB1*03:22 | HLA-DQA1*06:01-DQB1*03:22 |
| HLA-DQA1*05:05-DQB1*02:02 | HLA-DQA1*01:05-DQB1*06:04 | HLA-DQA1*04:01-DQB1*04:01 | HLA-DQA1*06:01-DQB1*04:01 |
| HLA-DQA1*05:05-DQB1*03:01 | HLA-DQA1*02:01-DQB1*03:87 | HLA-DQA1*05:01-DQB1*02:02 | HLA-DQA1*06:01-DQB1*04:02 |
| HLA-DQA1*05:05-DQB1*03:02 | HLA-DQA1*03:01-DQB1*02:01 | HLA-DQA1*05:01-DQB1*03:02 |  |

**Supplementary table 8:**
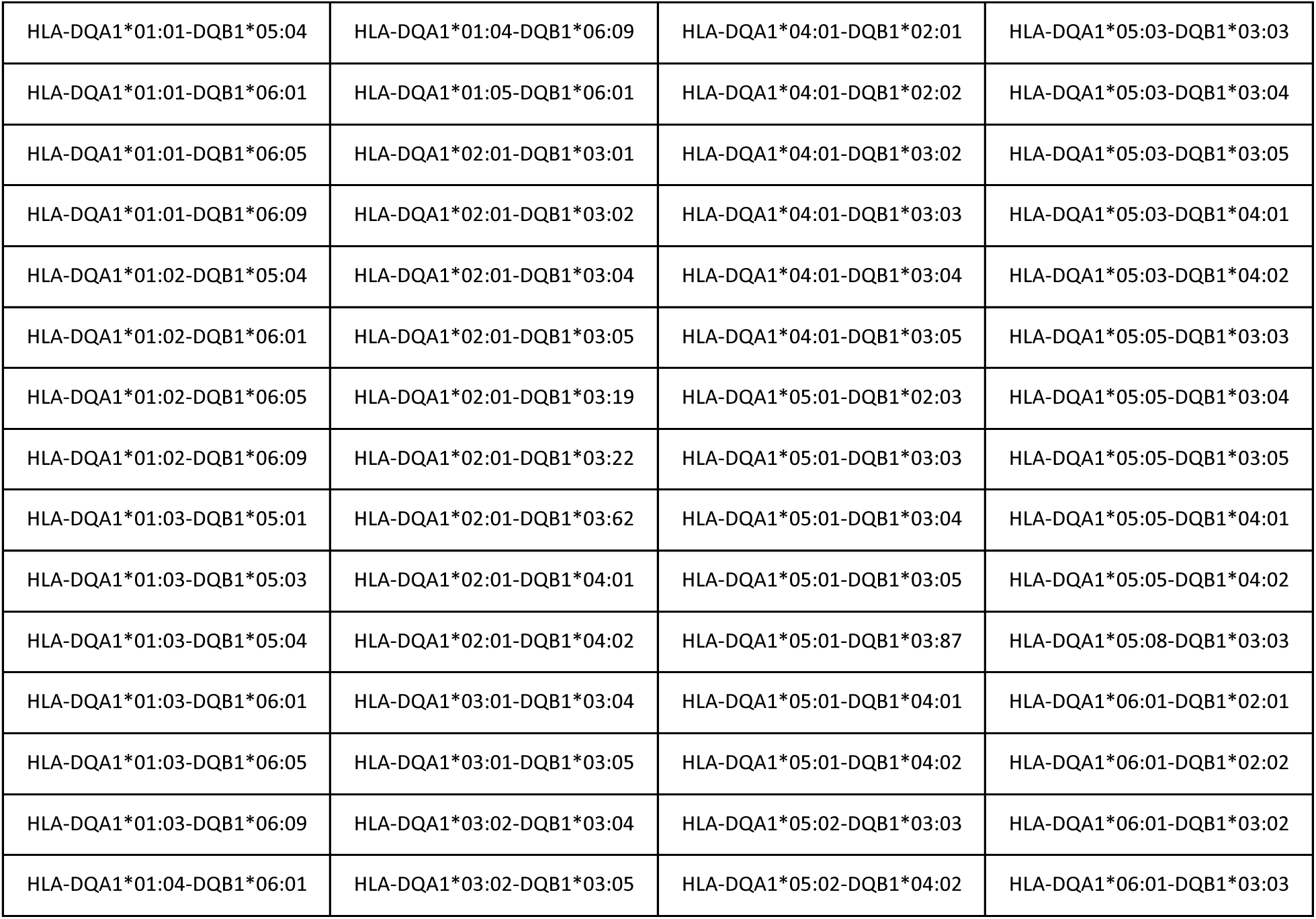
Prevalent DQ molecules with distance greater than 0.025 to molecules covered by the method including the novel data.

|  |  |  |  |
| --- | --- | --- | --- |
| HLA-DQA1*01:01-DQB1*05:04 | HLA-DQA1*01:04-DQB1*06:09 | HLA-DQA1*04:01-DQB1*02:01 | HLA-DQA1*05:03-DQB1*03:03 |
| HLA-DQA1*01:01-DQB1*06:01 | HLA-DQA1*01:05-DQB1*06:01 | HLA-DQA1*04:01-DQB1*02:02 | HLA-DQA1*05:03-DQB1*03:04 |
| HLA-DQA1*01:01-DQB1*06:05 | HLA-DQA1*02:01-DQB1*03:01 | HLA-DQA1*04:01-DQB1*03:02 | HLA-DQA1*05:03-DQB1*03:05 |
| HLA-DQA1*01:01-DQB1*06:09 | HLA-DQA1*02:01-DQB1*03:02 | HLA-DQA1*04:01-DQB1*03:03 | HLA-DQA1*05:03-DQB1*04:01 |
| HLA-DQA1*01:02-DQB1*05:04 | HLA-DQA1*02:01-DQB1*03:04 | HLA-DQA1*04:01-DQB1*03:04 | HLA-DQA1*05:03-DQB1*04:02 |
| HLA-DQA1*01:02-DQB1*06:01 | HLA-DQA1*02:01-DQB1*03:05 | HLA-DQA1*04:01-DQB1*03:05 | HLA-DQA1*05:05-DQB1*03:03 |
| HLA-DQA1*01:02-DQB1*06:05 | HLA-DQA1*02:01-DQB1*03:19 | HLA-DQA1*05:01-DQB1*02:03 | HLA-DQA1*05:05-DQB1*03:04 |
| HLA-DQA1*01:02-DQB1*06:09 | HLA-DQA1*02:01-DQB1*03:22 | HLA-DQA1*05:01-DQB1*03:03 | HLA-DQA1*05:05-DQB1*03:05 |
| HLA-DQA1*01:03-DQB1*05:01 | HLA-DQA1*02:01-DQB1*03:62 | HLA-DQA1*05:01-DQB1*03:04 | HLA-DQA1*05:05-DQB1*04:01 |
| HLA-DQA1*01:03-DQB1*05:03 | HLA-DQA1*02:01-DQB1*04:01 | HLA-DQA1*05:01-DQB1*03:05 | HLA-DQA1*05:05-DQB1*04:02 |
| HLA-DQA1*01:03-DQB1*05:04 | HLA-DQA1*02:01-DQB1*04:02 | HLA-DQA1*05:01-DQB1*03:87 | HLA-DQA1*05:08-DQB1*03:03 |
| HLA-DQA1*01:03-DQB1*06:01 | HLA-DQA1*03:01-DQB1*03:04 | HLA-DQA1*05:01-DQB1*04:01 | HLA-DQA1*06:01-DQB1*02:01 |
| HLA-DQA1*01:03-DQB1*06:05 | HLA-DQA1*03:01-DQB1*03:05 | HLA-DQA1*05:01-DQB1*04:02 | HLA-DQA1*06:01-DQB1*02:02 |
| HLA-DQA1*01:03-DQB1*06:09 | HLA-DQA1*03:02-DQB1*03:04 | HLA-DQA1*05:02-DQB1*03:03 | HLA-DQA1*06:01-DQB1*03:02 |
| HLA-DQA1*01:04-DQB1*06:01 | HLA-DQA1*03:02-DQB1*03:05 | HLA-DQA1*05:02-DQB1*04:02 | HLA-DQA1*06:01-DQB1*03:03 |

**Supplementary table 9:** Overview of donor cell lines and their HLA typing in the benchmark data from Marcu et al. 2021. For the donor Donor_BD_ZH14, the HLA typing was originally listed as having five DQ molecules. As four of these molecules shared two α- and two β-chains, the erroneous extra molecule was found to be DQA1*01:02-DQB1*02:01. This could be a misspelling of DPA1*01:02-DPB1*02:01, however, DPA1*01:02 has been removed from the IMGT/HLA database due to having an erroneous sequence (https://www.ebi.ac.uk/ipd/imgt/hla/alleles/allele/?accession=HLA00498). As such, we decided to delete DQA1*01:02-DQB1*02:01 from the HLA typing of this donor.

|  |  |
| --- | --- |
| Donor_16_113 | DRB1*04:01,DRB1*07:01,DQA1*01:01-DQB1*03:01,DQA1*01:01-DQB1*03:03 |
| Donor_BD_ZH01 | DRB1*07:01,DRB1*11:01,DRB3*02:02,DRB4*01:01,DQA1*02:01-DQB1*02:02,DQA1*05:01-DQB1*02:02,DPA1*01:03-DPB1*02:01,DPA1*01:03-DPB1*19:01 |
| Donor_BD_ZH02 | DRB1*04:01,DRB4*01:01,DQA1*03:01-DQB1*03:01,DQA1*03:02-DQB1*03:01,DPA1*01:03-DPB1*04:01,DPA1*01:03-DPB1*04:02 |
| Donor_BD_ZH03 | DRB1*07:01,DRB1*11:03,DRB3*02:02,DRB4*01:01,DQA1*02:01-DQB1*02:02,DQA1*02:01-DQB1*03:01,DQA1*05:05-DQB1*02:02,DQA1*05:05-DQB1*03:01,DPA1*01:03-DPB1*03:01,DPA1*01:03-DPB1*04:01 |
| Donor_BD_ZH04 | DRB1*07:01,DRB1*13:01,DRB3*01:01,DRB4*01:01,DQA1*01:03-DQB1*02:02,DQA1*01:03-DQB1*06:03,DQA1*02:01-DQB1*02:02,DQA1*02:01-DQB1*06:03,DPA1*01:03-DPB1*03:01,DPA1*01:03-DPB1*04:01,DPA1*02:02-DPB1*03:01,DPA1*02:02-DPB1*04:01 |
| Donor_BD_ZH05 | DRB1*11:01,DRB1*14:54,DRB3*02:02,DQA1*01:01-DQB1*03:01,DQA1*01:01-DQB1*05:03,DQA1*05:05-DQB1*03:01,DQA1*05:05-DQB1*05:03,DPA1*01:03-DPB1*04:01 |
| Donor_BD_ZH06 | DRB1*08:01,DRB1*13:03,DRB3*01:01,DQA1*04:01-DQB1*03:01,DQA1*04:01-DQB1*04:02,DQA1*05:05-DQB1*03:01,DQA1*05:05-DQB1*04:02,DPA1*01:03-DPB1*04:01,DPA1*01:03-DPB1*04:02 |
| Donor_BD_ZH07 | DRB1*04:01,DRB1*15:01,DRB4*01:01,DRB5*01:01,DQA1*01:02-DQB1*03:01,DQA1*01:02-DQB1*06:02,DQA1*03:01-DQB1*03:01,DQA1*03:01-DQB1*06:02,DPA1*01:03-DPB1*04:02,DPA1*01:03-DPB1*05:01 |
| Donor_BD_ZH08 | DRB1*13:03,DRB1*14:01,DQA1*05:05-DQB1*03:01 |
| Donor_BD_ZH09 | DRB1*07:01,DRB4*01:03,DQA1*02:01-DQB1*02:02,DPA1*01:03-DPB1*03:01,DPA1*01:03-DPB1*04:02 |
| Donor_BD_ZH11 | DRB1*07:01,DRB1*13:02,DRB3*03:01,DRB4*01:03,DQA1*01:02-DQB1*03:03,DQA1*01:02-DQB1*06:04,DQA1*02:01-DQB1*03:03,DQA1*02:01-DQB1*06:04,DPA1*01:03-DPB1*03:01,DPA1*01:03-DPB1*04:01 |
| Donor_BD_ZH12 | DRB1*01:01,DRB1*12:01,DRB3*02:02,DQA1*01:01-DQB1*03:01,DQA1*01:01-DQB1*05:01,DQA1*05:05-DQB1*03:01,DQA1*05:05-DQB1*05:01,DPA1*01:03-DPB1*02:01,DPA1*01:03-DPB1*09:01,DPA1*02:01-DPB1*02:01,DPA1*02:01-DPB1*09:01 |
| Donor_BD_ZH13 | DRB1*10:01,DRB1*15:01,DRB5*01:01,DQA1*01:01-DQB1*05:01,DQA1*01:01-DQB1*06:02,DQA1*01:02-DQB1*05:01,DQA1*01:02-DQB1*06:02,DPA1*01:03-DPB1*02:01,DPA1*01:03-DPB1*04:02 |
| Donor_BD_ZH14 | DRB1*04:01,DRB1*13:03,DRB3*01:01,DRB4*01:03,DQA1*03:02-DQB1*03:01,DQA1*03:02-DQB1*03:02,DQA1*05:05-DQB1*03:01,DQA1*05:05-DQB1*03:02,DPA1*01:03-DPB1*02:01 |
| Donor_BD_ZH15 | DRB1*03:01,DRB1*11:01,DRB3*01:01,DRB3*02:02,DQA1*05:01-DQB1*02:01,DQA1*05:01-DQB1*03:01,DPA1*01:03-DPB1*01:01,DPA1*01:03-DPB1*03:01,DPA1*02:01-DPB1*01:01,DPA1*02:01-DPB1*03:01 |
| Donor_BD_ZH16 | DRB1*03:01,DRB1*04:05,DQA1*03:02-DQB1*02:01,DQA1*03:02-DQB1*02:02,DQA1*05:01-DQB1*02:01,DQA1*05:01-DQB1*02:02 |
| Donor_BD_ZH17 | DRB1*14:54,DRB1*15:01,DRB3*02:02,DRB5*01:01,DQA1*01:01-DQB1*05:03,DQA1*01:01-DQB1*06:02,DQA1*01:02-DQB1*05:03,DQA1*01:02-DQB1*06:02,DPA1*01:03-DPB1*02:01 |

## Notes

### Competing Interest Statement

The authors have declared no competing interest.

### Summary of Updates

The revised version of the manuscript includes more background on the issue of cis and trans-only DQ molecules, as well as an external benchmark on DQ-heterozygous samples.

## References

1. Rocha, N. & Neefjes, J. MHC class II molecules on the move for successful antigen presentation. Embo Journal 27, 1–5 (2008).

2. Reynisson, B. et al. Improved Prediction of MHC II Antigen Presentation through Integration and Motif Deconvolution of Mass Spectrometry MHC Eluted Ligand Data. J Proteome Res 19, 2304–2315 (2020).

3. Arango, M. T. et al. HLA-DRB1 the notorious gene in the mosaic of autoimmunity. Immunol Res 65, 82–98 (2017).

4. Erlich, H. et al. HLA DR-DQ haplotypes and genotypes and type 1 diabetes risk analysis of the type 1 diabetes genetics consortium families. Diabetes 57, 1084–1092 (2008).

5. Hu, X. et al. Additive and interaction effects at three amino acid positions in HLA-DQ and HLA-DR molecules drive type 1 diabetes risk. Nat Genet 47, 898–905 (2015).

6. Stepniak, D. et al. Large-scale characterization of natural ligands explains the unique gluten-binding properties of HLA-DQ2. Journal of Immunology 180, 3268–3278 (2008).

7. Racle, J. et al. Machine learning predictions of MHC-II specificities reveal alternative binding mode of class II epitopes. bioRxiv 2022.06.26.497561 (2022) doi:10.1101/2022.06.26.497561.

8. Bergseng, E. et al. Different binding motifs of the celiac disease-associated HLA molecules DQ2.5, DQ2.2, and DQ7.5 revealed by relative quantitative proteomics of endogenous peptide repertoires. Immunogenetics 67, 73–84 (2014).

9. Sidney, J. et al. Divergent motifs but overlapping binding repertoires of six HLA-DQ molecules frequently expressed in the worldwide human population. Journal of Immunology 185, 4189–4198 (2010).

10. Vartdal, F. et al. The peptide binding motif of the disease associated HLA-DQ (α 1* 0501, β 1* 0201) molecule. Eur J Immunol 26, 2764–2772 (1996).

11. Tollefsen, S. et al. Structural and functional studies of trans-encoded HLA-DQ2.3 (DQA1*03:01/DQB1*02:01) protein molecule. Journal of Biological Chemistry 287, 13611–13619 (2012).

12. Kwok, W. W., Kovats, S., Thurtle, P. & Nepom, G. T. HLA-DQ allelic polymorphisms constrain patterns of class II heterodimer formation. The Journal of Immunology 150, 2263–2272 (1993).

13. Creary, L. E. et al. High-resolution HLA allele and haplotype frequencies in several unrelated populations determined by next generation sequencing: 17th International HLA and Immunogenetics Workshop joint report. Hum Immunol 82, 505–522 (2021).

14. Petersdorf, E. W. et al. HLA-DQ heterodimers in hematopoietic cell transplantation. Blood 139, 3009–3017 (2022).

15. Lundin, K. E. et al. T lymphocyte recognition of a celiac disease-associated cis- or trans-encoded HLA-DQ alpha/beta-heterodimer. The Journal of Immunology 145, 136–139 (1990).

16. Kwok, W. W. & Nepom, G. T. Structural and functional constraints on HLA class II dimers implicated in susceptibility to insulin dependent diabetes mellitus. Baillieres Clin Endocrinol Metab 5, 375–393 (1991).

17. McFarland, B. J. & Beeson, C. Binding interactions between peptides and proteins of the class II Major Histocompatibility Complex. Med Res Rev 22, 168–203 (2002).

18. Nielsen, M., Andreatta, M., Peters, B. & Buus, S. Immunoinformatics: Predicting Peptide– MHC Binding. Annu Rev Biomed Data Sci 3, 191–215 (2020).

19. Reynisson, B., Alvarez, B., Paul, S., Peters, B. & Nielsen, M. NetMHCpan-4.1 and NetMHCIIpan-4.0: improved predictions of MHC antigen presentation by concurrent motif deconvolution and integration of MS MHC eluted ligand data. Nucleic Acids Res 48, W449–W454 (2020).

20. Gfeller, D. & Bassani-Sternberg, M. Predicting antigen presentation-What could we learn from a million peptides? Front Immunol 9, 1716 (2018).

21. Nielsen, M., Lund, O., Buus, S. & Lundegaard, C. MHC Class II epitope predictive algorithms. Immunology 130, 319–328 (2010).

22. Bassani-Sternberg, M. et al. Direct identification of clinically relevant neoepitopes presented on native human melanoma tissue by mass spectrometry. Nat Commun 7, 13404 (2016).

23. Kaabinejadian, S. et al. Accurate MHC Motif Deconvolution of Immunopeptidomics Data Reveals a Significant Contribution of DRB3, 4 and 5 to the Total DR Immunopeptidome. Front Immunol 13, 835454 (2022).

24. Alvarez, B., Barra, C., Nielsen, M. & Andreatta, M. Computational Tools for the Identification and Interpretation of Sequence Motifs in Immunopeptidomes. Proteomics 18, 1700252 (2018).

25. Caron, E. et al. Analysis of major histocompatibility complex (MHC) immunopeptidomes using mass spectrometry. Molecular and Cellular Proteomics 14, 3105–3117 (2015).

26. Purcell, A. W., Ramarathinam, S. H. & Ternette, N. Mass spectrometry–based identification of MHC-bound peptides for immunopeptidomics. Nat Protoc 14, 1687–1707 (2019).

27. Barra, C. et al. Footprints of antigen processing boost MHC class II natural ligand predictions. Genome Med 10, (2018).

28. Paul, S. et al. Determination of a predictive cleavage motif for eluted major histocompatibility complex class II ligands. Front Immunol 9, 1795 (2018).

29. Racle, J. et al. Robust prediction of HLA class II epitopes by deep motif deconvolution of immunopeptidomes. Nat Biotechnol 37, 1283–1286 (2019).

30. Wang, P. et al. Peptide binding predictions for HLA DR, DP and DQ molecules. BMC Bioinformatics 11, 568 (2010).

31. Nielsen, M., Lundegaard, C. & Lund, O. Prediction of MHC class II binding affinity using SMM-align, a novel stabilization matrix alignment method. BMC Bioinformatics 8, 238 (2007).

32. Alvarez, B. et al. NNAlign_MA; MHC peptidome Deconvolution for Accurate MHC Binding Motif Characterization and Improved T-cell Epitope Predictions. Molecular and Cellular Proteomics 18, 2459–2477 (2019).

33. Nielsen, M. & Andreatta, M. NetMHCpan-3.0; improved prediction of binding to MHC class I molecules integrating information from multiple receptor and peptide length datasets. Genome Med 8, (2016).

34. Hoof, I. et al. NetMHCpan, a method for MHC class I binding prediction beyond humans. Immunogenetics 61, 1–13 (2009).

35. Karosiene, E. et al. NetMHCIIpan-3.0, a common pan-specific MHC class II prediction method including all three human MHC class II isotypes, HLA-DR, HLA-DP and HLA-DQ. Immunogenetics 65, 711–724 (2013).

36. Gonzalez-Galarza, F. F., Christmas, S., Middleton, D. & Jones, A. R. Allele frequency net: A database and online repository for immune gene frequencies in worldwide populations. Nucleic Acids Res 39, (2011).

37. Thomsen, M. C. F., Lundegaard, C., Buus, S., Lund, O. & Nielsen, M. MHCcluster, a method for functional clustering of MHC molecules. Immunogenetics 65, 655–665 (2013).

38. Moore, R. M., Harrison, A. O., McAllister, S. M., Polson, S. W. & Eric Wommack, K. Iroki: Automatic customization and visualization of phylogenetic trees. PeerJ 8, e8584 (2020).

39. Larkin, M. A. et al. Clustal W and Clustal X version 2.0. Bioinformatics 23, 2947–2948 (2007).

40. Marcu, A. et al. HLA Ligand Atlas: A benign reference of HLA-presented peptides to improve T-cell-based cancer immunotherapy. J Immunother Cancer 9, (2021).

